# Stress-Induced Alteration of Small Extracellular Vesicles Drives Amyloid-Beta Sequestration and Exacerbates Alzheimer’s Disease Pathogenesis

**DOI:** 10.1101/2025.05.23.655679

**Authors:** Sadaqa Zehra, Sanskriti Rai, Komal Rani, Saumitra Dey Choudhury, Himanshu Rai, Suchismita Bhowmik, Nitin Mohan, Anu Gupta, Prasun Chatterjee, Thota Jagadeshwar Reddy, Neerja Rani, Gyan Prakash Modi, Fredrik Nikolajeff, Saroj Kumar

## Abstract

While small extracellular vesicles (sEVs) are implicated in amyloid-beta (Aβ) trafficking, the mechanisms governing their interaction with Aβ aggregates and plaque formation remain unresolved. Here, we report a paradigm-shifting discovery: sEVs undergo dynamic structural remodelling in response to stress, enabling selective binding to Aβ aggregates-a phenomenon absent under normal physiological conditions. Using multimodal stressors, including mechanical (ultrasonication/agitation), physical (hyperthermia), and biological (oxidative damage), we demonstrate that stress-modified sEVs exhibit high-affinity binding to small Aβ aggregates (SA) through scaffold reorganization, as validated by super-resolution microscopy and quantitative colocalization assays. Crucially, these remodelled sEVs act as potent carriers, enhancing SA internalization by neuronal cells *in vitro*. Strikingly, in post-mortem Alzheimer’s disease (AD) brains and APP-PS1 transgenic mice, sEVs were spatially enriched at amyloid plaque margins, suggesting a direct role in Aβ sequestration and plaque expansion. Consistent with clinical relevance, sEVs isolated from AD patients exhibited an intrinsic SA-binding capacity, recapitulating stress-induced interactions observed experimentally. Our findings reveal that stress-primed sEVs function as pathological chaperones, binding to and internalizing Aβ aggregates, thereby accelerating plaque nucleation and disease progression. This study provides the first evidence of stress-mediated sEV plasticity as a critical driver of Aβ pathology, redefining therapeutic strategies targeting extracellular vesicle biology in neurodegenerative disorders.

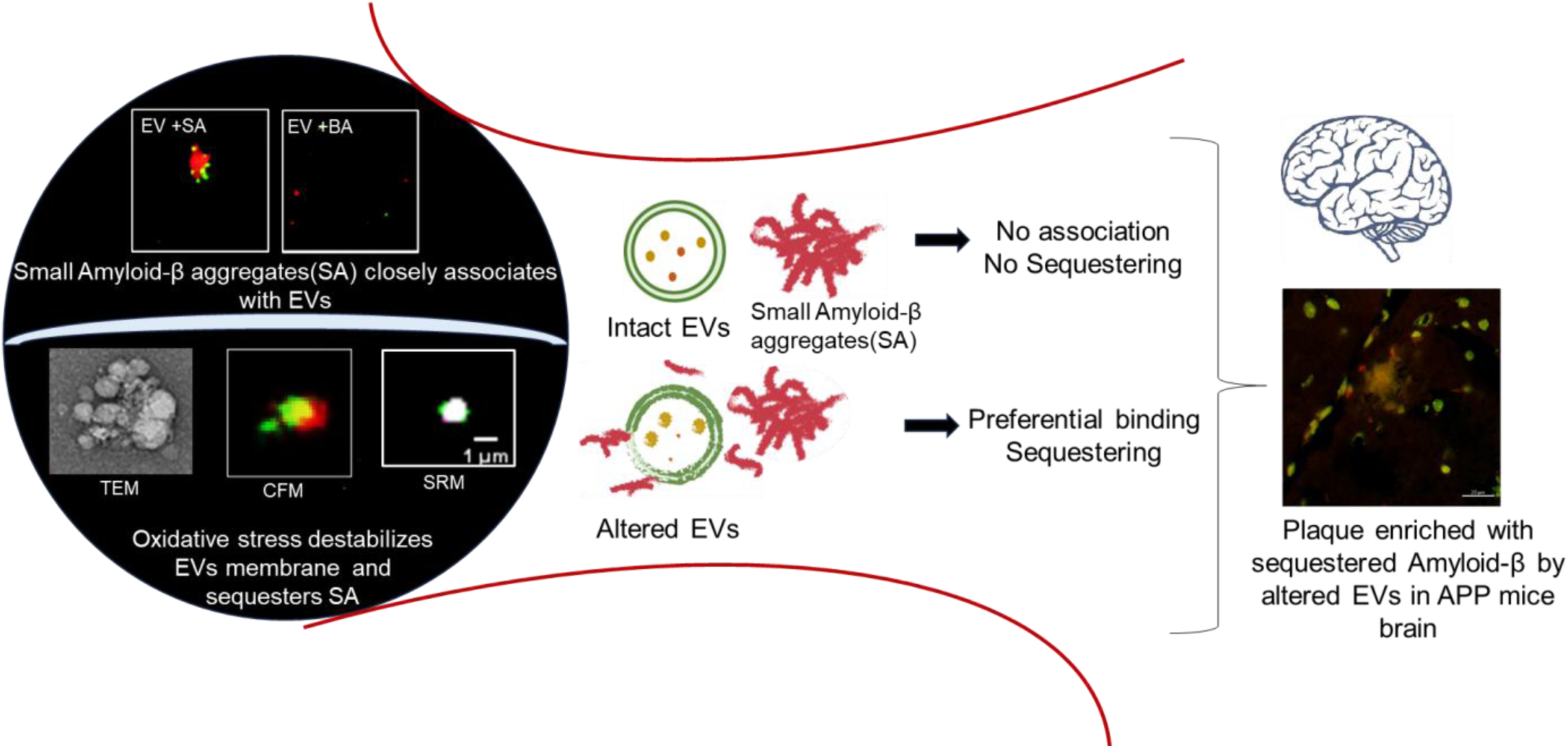

## 1. Introduction

Alzheimer’s disease (AD) is the most prevalent form of dementia, associated with an aging population, and is a major worldwide health problem (1). The underlying neuropathology of AD is strongly attributed to the accumulation of intracellular neurofibrillary tangles made of hyperphosphorylated tau protein and extracellular amyloid-beta (Aβ) plaques (2). Aβ peptides, particularly Aβ42, have been reported to be prone to misfolding and aggregation (3).

Small extracellular vesicles (sEVs) are nano-sized, membrane-bound particles that are secreted by virtually all cell types into the extracellular space (4). Typically ranging from 30 to 150 nanometers in diameter, sEVs have been extensively studied in recent years due to their essential role in intercellular communication (5). The specific molecular content of sEVs comprising proteins, lipids, and various RNA species, including mRNAs and microRNAs, is dictated by the originating cell type and its physiological or pathological state and has profound relevance in different physiological processes and pathological conditions, including neurodegenerative diseases like AD (6). The sEVs have been shown to carry amyloid-beta (Aβ) and tau proteins, key pathological markers of the disease, facilitating their spread across different brain regions and contributing to the progressive nature of neurodegeneration observed in Alzheimer’s disease (7,8). The sEVs also function as Aβ scavengers by ensnaring Aβ through the glycosphingolipids on their surface (9). In-depth electron microscopy analyses of the brains of transgenic APP mice have revealed the presence of neuronal endosomes that contain Aβ, notably MVBs (10). Recent studies have shown that extracellular plaques primarily arise from intraneuronal β-amyloid accumulation within membrane tubules, forming a central amyloid “core” which then degenerates to produce classical senile plaques (11). In 2006, Rajendran et al. observed that APP-transfected cells released small amounts of Aβ in association with exosomes and they also reported the exosomal marker Alix to be around human amyloid plaques, suggesting that exosome-associated Aβ plays a role in plaque formation (12). In the context of exosomes, it is presumed that Aβ is topologically associated with the surface membrane (13). Immuno-electron microscopy analysis of exosomes isolated from N2a cells expressing APP revealed surface-bound Aβ (14). Research into the molecular mechanism underlying the binding of Aβ to exosomes has been ongoing. Recent findings indicate that Aβ interacts with glycosphingolipids (GSLs), which tend to collect on the membrane’s outer layer, exposing their glycans to the outside environment. Aβ recognizes and binds to these GSL clusters (15).

There is heterogeneity in scientific studies discussing the association of EVs with amyloid-β, where some groups have reported the role of EVs in reducing the rate of Aβ (1–42) fibril formation by posing interference in the fibril elongation step (16). In addition to transporting Aβ and tau, sEVs in AD are thought to participate in the neuro-inflammatory response by transporting pro-inflammatory molecules, sEVs activate microglia and astrocytes, exacerbating neuronal damage (17,18).

Alzheimer’s disease progression is attributed to multiple pathophysiologic mechanisms (19). Studies have discussed the role of oxidative stress on AD progression and numerous antioxidant drugs have been in trial for AD therapeutics (20–23). Oxidative stress increases EV production by promoting MVB degradation (24) and alters EV cargo composition, affecting oxidative states in recipient cells (25). It also contributes to β-sheet-rich fibril formation of Aβ42 (26). Additionally, oxidative stress oxidizes amyloid-β into toxic oligomers and alters EV lipid bilayer conformation through lipid peroxidation, driving disease progression (27,28). We have studied the EV-Aβ association under mechanical, physical, and biological stress conditions, explaining how oxidative stress acts as a disease-relevant pathophysiological process that alters EV scaffold and enhances their affinity for Aβ.

## 2. Results

### 2.1. Characterization and validation of isolated EVs

Plasma-derived sEVs were successfully isolated, characterized, and validated. In Figure 1, the plasma-derived sEVs were morphologically characterized by transmission electron microscopy where spherical lipid bi-layered vesicles were observed in the size range of 65-80 nm in diameter (Figure 1A, Supplementary Figure 1A). PsEVs were also visualized by Confocal Microscopy using Alexa Fluor 488 tagged anti-CD9 antibodies (Figure 1B, Supplementary Figure 1B-C). The size distributions and concentration of plasma-derived sEVs were observed by nanoparticle tracking analysis. The mean concentration of plasma-derived sEVs was 5.0E+10 particle/ml (Figure 1C). Western blot validated the presence of sEVs-specific markers (surface marker: CD9, CD63, CD81, and luminal marker: TSG101) in the isolated sEVs (Figure 1D, Supplementary Figure 1D). We also assessed the level of co-isolated contaminating protein like apolipoprotein in our sample (F3-F6) for evaluating sample purity (Figure 1D).

**Figure 1:**
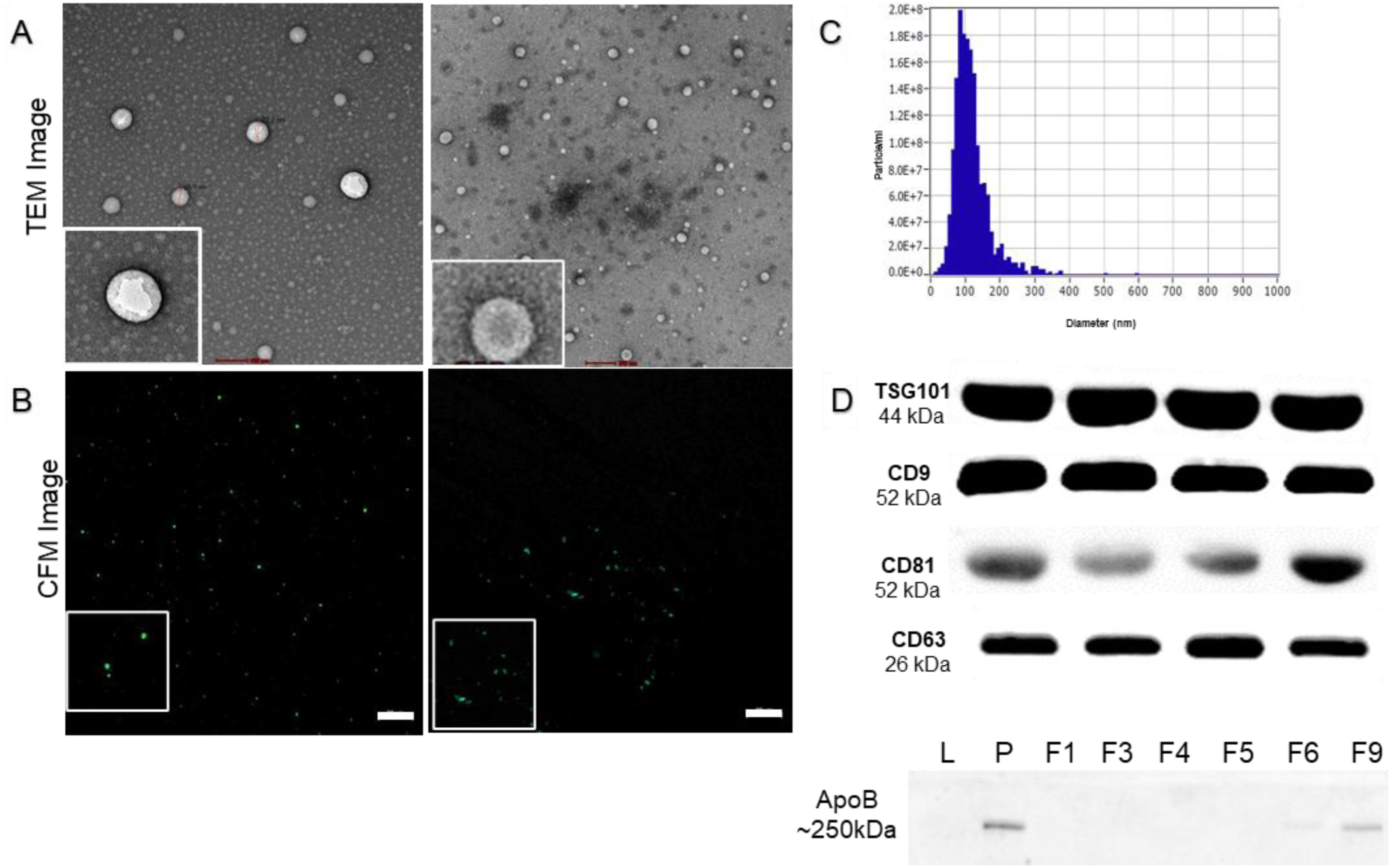
Characterization of isolated EVs through transmission electron microscopy (A) and confocal microscopy (B). Nanoparticle Analysis (NTA) showing the size distribution of EVs (nm) and particle concentration (particle/ml) (C); Western blot of EVs markers (CD9, CD63, CD81, Tsg101) and protein co-isolate (Apolipoprotein-B) (D). Scale bar= 20µm

### 2.2. Characterization of different Amyloid-β aggregates

To mimic different amyloid-β species relevant to AD pathology, we prepared different-sized aggregates of Aβ-42 and confirmed their structural morphology and size distribution, through transmission electron microscopy, confocal microscopy, dynamic light scattering analysis, nanoparticle tracking analysis and Thioflavin-T (ThT) fluorescence. Lyophilized Amyloid-β was resuspended in 60mM NaOH with pH adjusted to 7.4; this starting solution was termed unaggregated amyloid beta peptides (UA) and immediately stored to-80℃ until further use. Small amyloid beta aggregates (SA) were prepared by ultrasonication, and fibrillar aggregates or big aggregates (BA) were prepared by ultrasonication followed by mild agitation for an hour. TEM micrograph revealed the representative aggregated structures both in SA and BA samples. UA group did not form any large visible aggregates, while the SA group formed visible globular aggregates, and the BA group exhibited typical features of short-rod-like protofibrils (Figure 2A, Supplementary Figure 2A). Following this, we also incubated all three groups, i.e., UA, SA, and BA, with AlexaFluor647 Anti-Aβ 42 Antibodies and observed under 40X magnification in a Confocal Microscope (Figure 2B). The CFM images reveal the distinct variations in fluorescent signals between these groups, which complement TEM results and were specific to Amyloid-β (Supplementary figure 2B-C). A distinctive difference in size and conformation was observed between the groups. Finally, we also assessed the size distribution using DLS and NTA, which further supported our claims on the nomenclature of the prepared aggregates based on their size (Figure 2C). DLS findings revealed the median size for UA, SA and BA was approximately 9, 100 and 1000 nm respectively (Figure 2C). In NTA, additional peaks were observed in the three groups, which provided clearer insight into the size distribution of three groups viz; UA, SA and BA (Figure 2C). Finally, we assessed the ThT fluorescence for each BA, SA and UA groups and observed enhanced fluorescence in BA group as a result of ThT binding to the β-sheet in the fibrillar structure of BA (Supplementary figure 2D).

**Figure 2:**
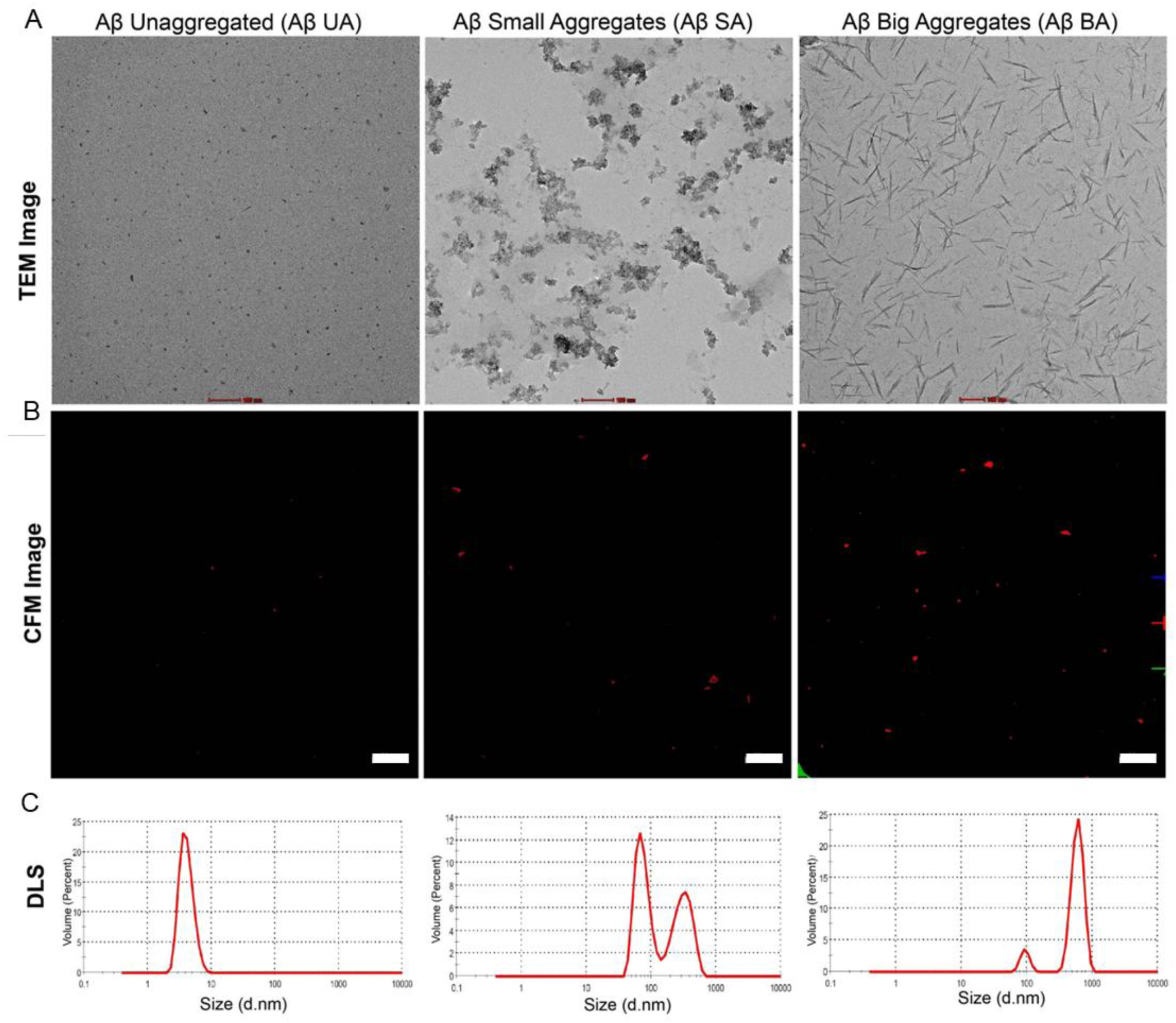
Characterization of different amyloid-β aggregate forms. Transmission electron micrographs (A) showed distinct morphology of the different Aβ aggregates. Confocal micrographs show increasing signal intensity according to Aβ aggregate size (B). Dynamic light scattering analysis (C) A= Unaggregated Aβ form (∼10nm); SA= Small Aβ aggregates (∼100nm); BA= Big Aβ aggregates (∼1000nm). Scale bar= 20µm.

### 2.3. Oxidative stress alters EVs’ driving amyloid-β-EVs association

Oxidative stress plays a well-established role in the pathogenesis of Alzheimer’s disease, as the brain exhibits oxidative damage resulting from excessive production of reactive oxygen species (ROS), which triggers oxidative stress and inflammatory responses, thereby accelerating AD progression. Hydrogen peroxide (H_2_O_2_) is considered a major ROS contributor and is implicated to play a key role in the pathogenesis of several progressive neurodegenerative diseases. Oxidative stress conditions induce substantial modifications in EV membrane composition, leading to altered biophysical properties that facilitate the sequestration and aggregation of pathogenic proteins such as amyloid-β (29). Therefore, in our study, H_2_O_2_-induced cytotoxicity was adopted as an oxidative stress model. EVs that were not exposed to any stress viz; (H_2_O_2_ and Ultrasonication) served as the control group (Figure 3.1 A). We subjected EVs to varying concentrations of H₂O₂ (100 µM, 10 µM, and 1 µM) for 24 hours where we observed that higher concentrations led to the formation of large aggregated masses (Supplementary Figure 3A). Therefore, we selected the minimal dose of 1 µM as a stress inducer. After incubating EVs with a 1µM H_2_O_2_ solution for 24 hours, we compared the H_2_O_2_-treated EVs with ultrasonicated EVs (Supplementary Figure 3B). TEM images revealed the structural morphology of EVs was distorted in H₂O₂-treated group compared to that of control EVs indicating alteration in EVs structural integrity. Similarly, CFM and SRM images also complemented TEM findings where by normal EVs do not form coaggregates with Aβ while H_2_O_2_-treated EVs associate with Aβ. These altered EVs showed self-aggregation, even at low temperatures (4℃) (Figure 3.1 B, Supplementary Figure 3C). dSTORM imaging also complemented TEM and SRM findings (Figure 3.1 C).

**Figure 3.1:**
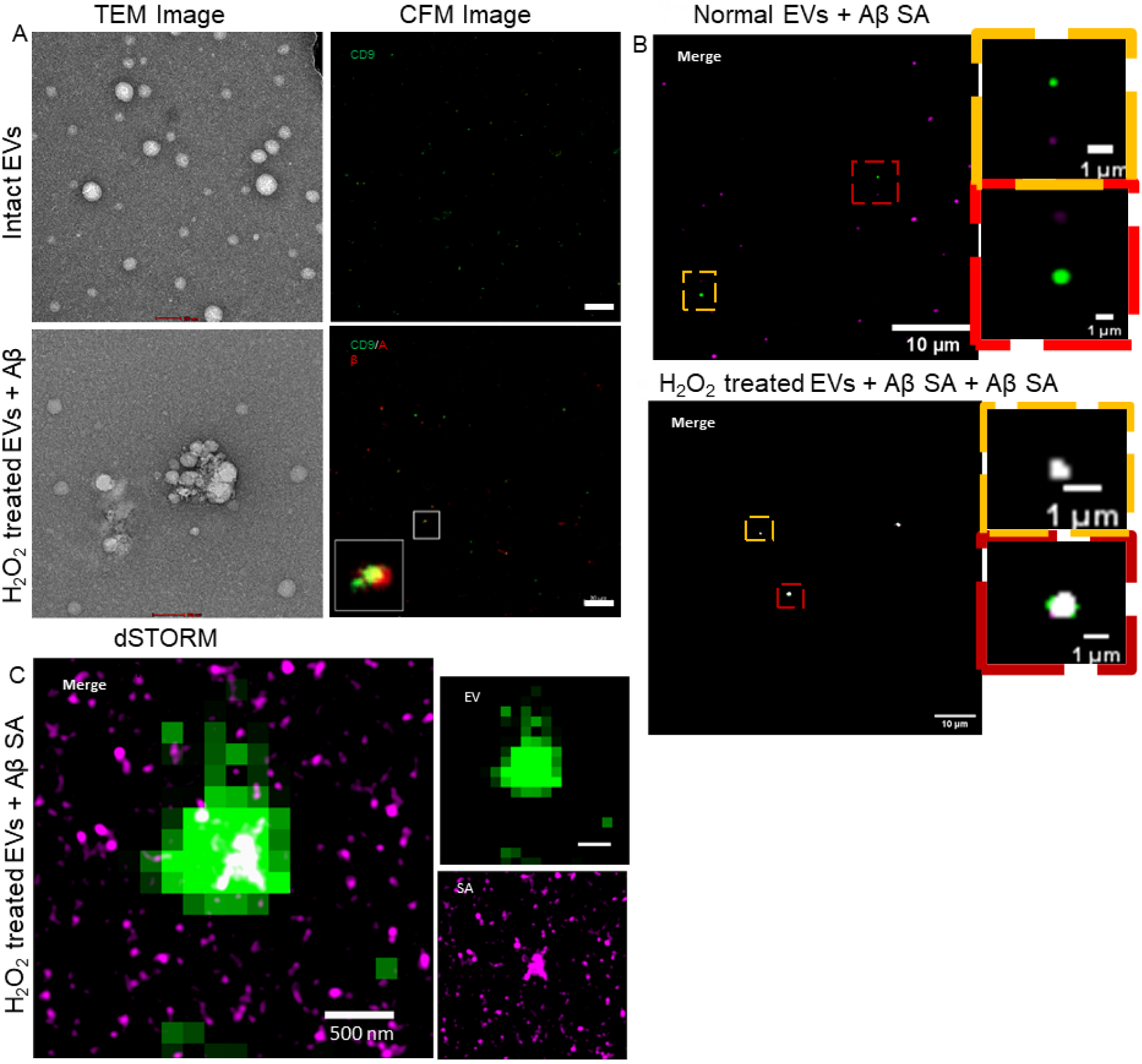
TEM and CFM micrograph shows aggregated structure in H_2_O_2_-treated EVs and Aβ aggregates (A). Comparison of interaction between normal and H_2_O_2_-treated EVs (B). The CFM and SRM micrographs also show signal overlap in H_2_O_2_-treated EVs only. Colour yellow is the merged signal of EVs (Green) and Aβ (red). Scale bar= 20µm. dSTORM Image shows EV-Aβ coaggregates (C). Colour white is the merged signal of EVs (Green) and Aβ (Magenta).

**Figure 3.2:**
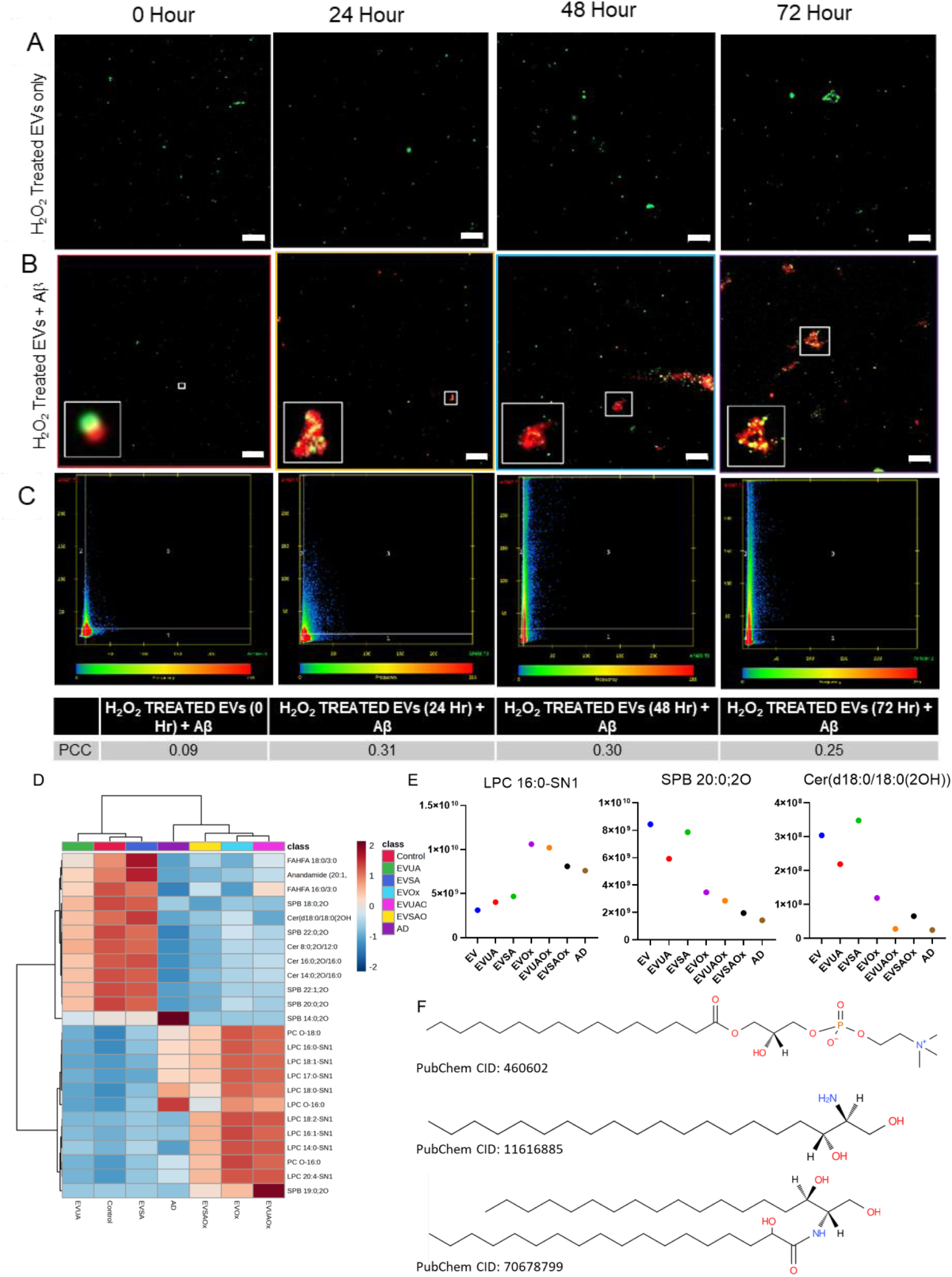
Temporal assessment of small amyloid-β aggregates and EV association at low temperature 4℃. An increase in H_2_O_2_-treated EV size was observed (A). Increased colocalization in EV-Aβ signals suggests an active sequestering by altered EVs in a time-dependent manner (B) and Colocalization coefficient (C). Colour yellow is the merged signal of EVs (Green) and Aβ (red). Scale bar= 20μm. Lipidomic analysis of EV lipids in different groups (D). Top dysregulated lipid group (E). Chemical structure of each lipid (F)

Furthermore, to monitor the effect of oxidative stress on EV-Aβ interaction, we subjected EVs to 1µM H_2_O_2_ stress for 24, 48 and 72 hours and incubated with Aβ SA for 2 hours at 4℃ (Figure 3.2 A-C). CFM images revealed an increase in EV signal intensity, indicating self-aggregation due to structural alterations in EVs, consistent with our TEM observations. We observed a gradual increase in colocalization of EV-Aβ signals, which indicated active sequestration by the altered EVs over time. An increase in EV size further supported these observations. (Supplementary Figure 3D). Pearson’s Correlation Coefficient was used to estimate the measure of colocalization for different time points which were; 0 hrs= 0.09, 24 hours= 0.31, 48 hours= 0.30 and 72hrs= 0.25) suggesting more association between altered EVs which plateaus after 72 hours.

Lipidomic analysis of AD EVs revealed elevated levels of Oxidized Glycerophospholipid (Supplementary Figure 3E), which forms the structural framework of EV membranes (30). A comparative analysis of the lipidome in Alzheimer’s disease (AD)-derived extracellular vesicles (EVs), normal EVs, and hypoxia-treated EVs with SA was conducted using untargeted lipidomic analysis (Figure 3D). This analysis revealed that lipid classes such as lysophosphatidylcholine (LPC), sphinganine (SPB), and ceramide (Cer) were highly dysregulated in stress-altered EVs compared to normal EVs and showed similar dysregulation patterns to those seen in AD-derived EVs (Figure 3E-F). Notably, lipid levels in the AD group closely mirrored those observed in EVs exposed to oxidative stress (Figure 3E). These results highlight the effects of oxidation on EV membrane composition, which further influence Aβ sequestration and aggregation.

### 2.4. Amyloid-β aggregates size regulate their association with EVs

After testing the association between EVs and Amyloid-β species with exposure to oxidative stresses, we further assessed whether this association depends on the size of Aβ itself. We prepared different-sized aggregates of Aβ-42, specifically assigned them as UA, SA, and BA following extensive characterization by TEM, CFM, DLS, and ThT assay (Figure 4, Supplementary Figure 4). Next, we incubated each group viz; UA, SA, and BA with freshly isolated EVs to evaluate the association. We examined if the different-sized aggregates affected their association with EVs using TEM and CFM. The TEM images revealed a characteristic cluster in the EVs and SA group, which was not present in either the UA or BA group suggesting EVs had a preferential association with the SA group (Figure 4A, Supplementary Figure 4A). TEM images of EVs with UA and BA showed no association between EVs and Aβ. Consequently, confocal microscopy confirmed that the SA has enhanced binding toward EVs (Figure 4B), as depicted by the colocalized signal (Yellow colour) from both EVs and Aβ-42 antibodies signals. CFM images did not reveal any colocalized signals for EVs with UA and BA groups (Supplementary Figure 4B). Co-localization analysis was used to estimate the measure of colocalization Pearson’s Correlation Coefficient for the EV+ SA group was 0.88 suggesting more association between EVs and SA compared to UA (PCC= 0.07) and BA group (PCC= 0.02) (Figure 4C). This supports the conclusion that SA preferentially binds to EVs compared to other amyloid-β aggregates.

**Figure 4:**
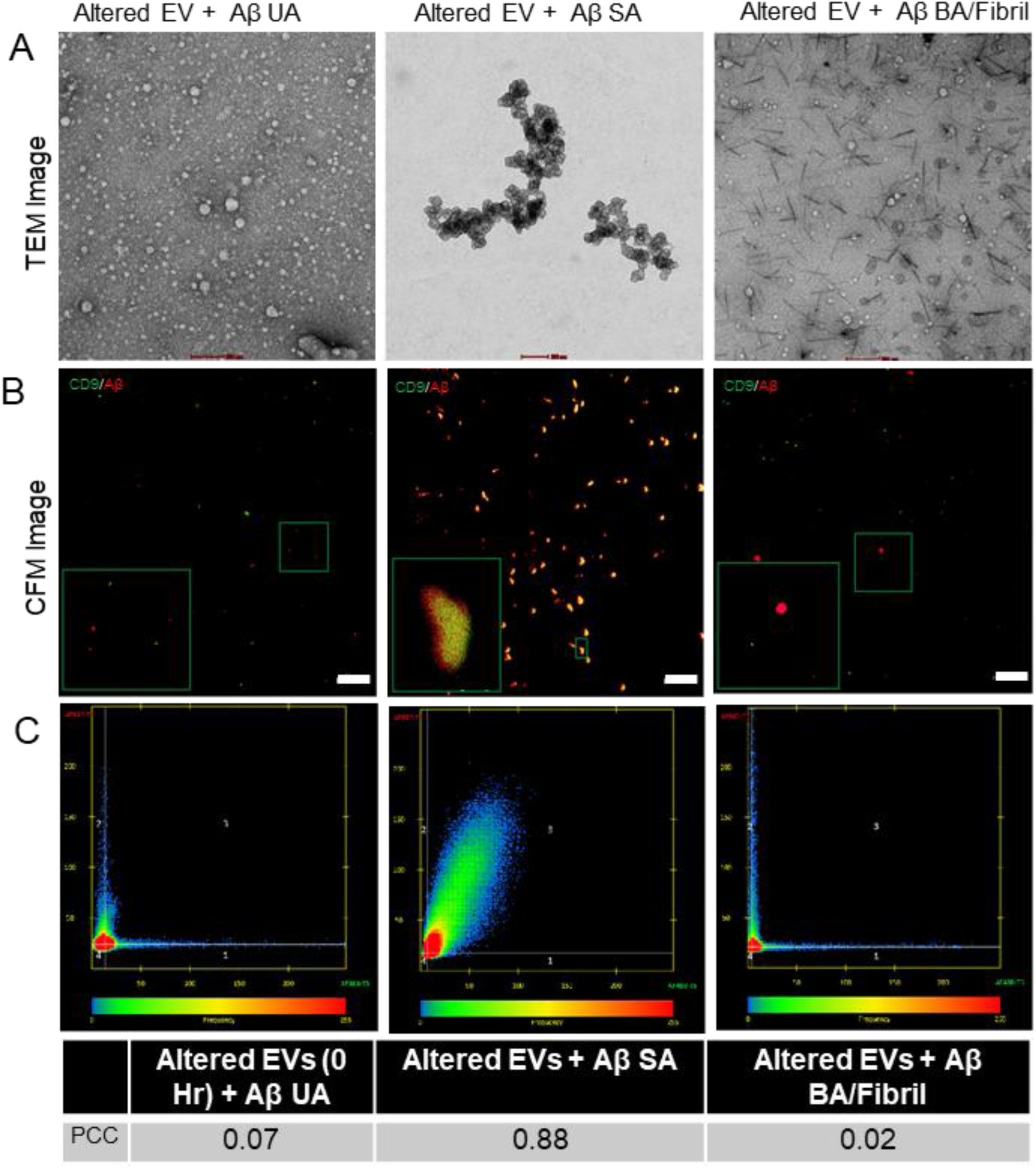
*In-vitro* assessment of different forms of amyloid-β and EV association. (Aβ42 UA= Unaggregated, Aβ42 SA= Small Aggregates, Aβ42 BA/Fibril= Big aggregates). TEM micrograph shows the distinct aggregated structures (A). Confocal microscopy shows enhanced SA binding to EVs (B), with a colocalization Pearson’s coefficient of 0.88, indicating stronger EV-SA association than with UA and BA (C). Colour yellow is the merged signal of EVs (Green) and Aβ (red). Scale bar= 20μm.

### 2.5. Assessment of EVs-Aβ association in a time-dependent manner

There is an existing ambiguity regarding the effect of EVs on extracellular amyloid-β. In line with our observations of significant findings pertaining to EVs-Aβ association as assessed by TEM and Confocal microscopy, we proceeded with studying the EVs and amyloid-β incubated together at different time intervals viz, 24, 48, and 72 hours at physiological temperature of 37℃ and low temperature 4℃ as control. We observed a gradual increase in the size of amyloid-β aggregates at 37℃, indicating active oligomerization over time (Figure 5A). Similarly, we also noted an increase in EV signal intensities, possibly related to EV self-aggregation at 37 ℃ (Supplementary Figure 5A). Finally, the EV-Aβ association also increased over time, as shown by the PCC values: 0.47 at 24 hours, 0.82 at 48 hours, and 0.90 at 72 hours, all at 37℃ (Figure 5A). Additionally, we also repeated the same experiment at 4℃ (Figure 5B), where low signal intensity was observed in both EVs and amyloid-β. It is well known that the storage conditions significantly influence the integrity and functionality of extracellular vesicles (EVs) (31,32). At 4℃, no close association was observed when EVs and Aβ were incubated together as indicated by PCC values of 0.0 at 24 hours, 0.15 at 48 hours, and 0.14 at 72 hours (Figure 5B). This suggests that the higher signal intensities at 37°C were due to amyloid-β sequestration mediated by EV-fragmentation (Supplementary Figure 5B). These findings imply that the observed association may reflect active Aβ oligomerization and EV membrane conformational changes under higher temperature conditions as primary mediators behind the close association between EVs and Aβ. Additionally, EV aggregation appeared to contribute to Aβ sequestration, particularly in the 37°C experimental group. Finally, we concluded that EVs membrane integrity is affected by temperature and influences its association with amyloid-β aggregates. EVs aggregation may promote amyloid-β aggregation, potentially making it a pathophysiological factor relevant to AD pathology.

**Figure 5:**
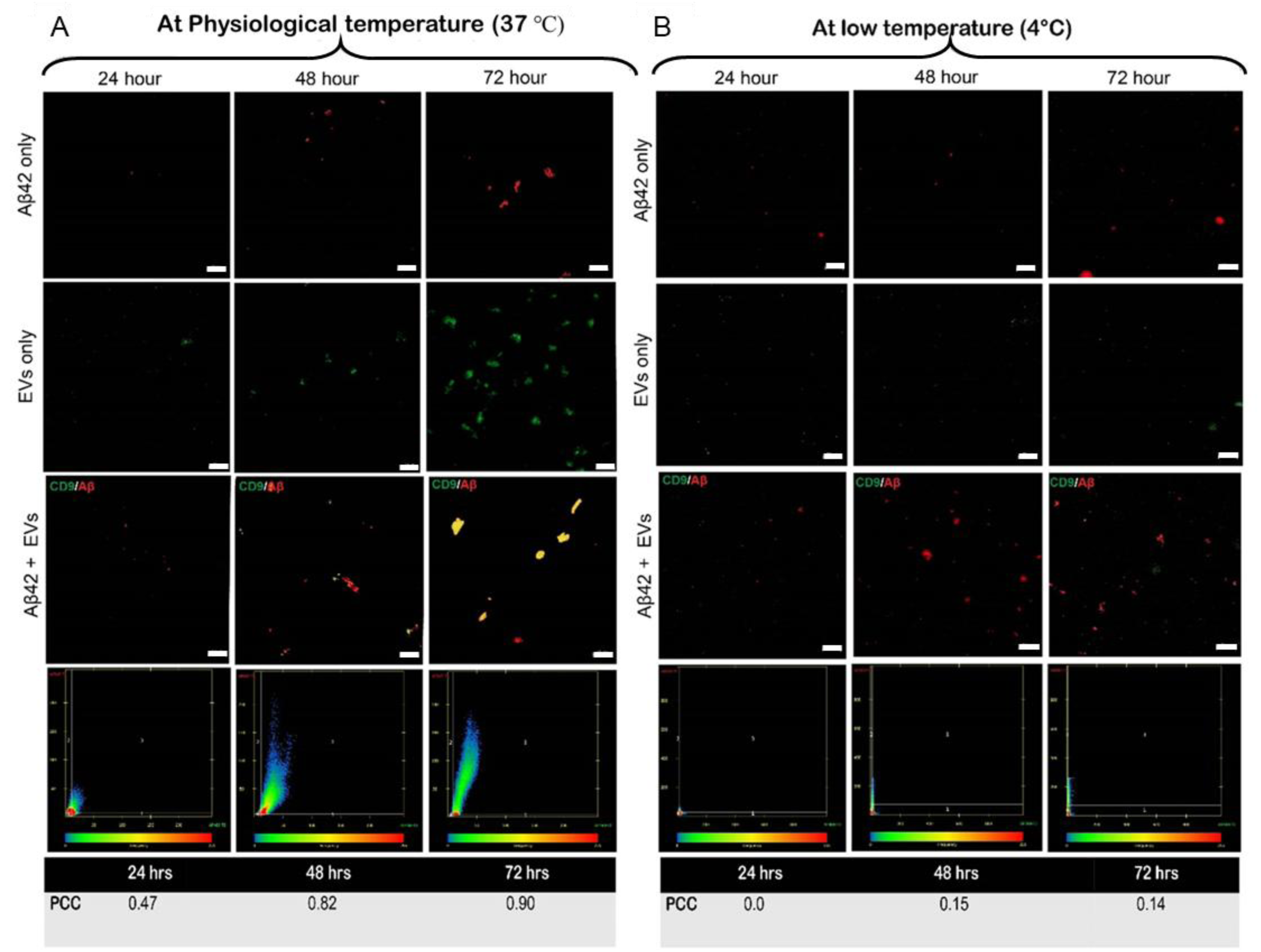
Temporal assessment of different forms of amyloid-β and EV association: at physiological temperature 37℃ (A)-Colour (yellow) represents the merged image of EVs (Green) and Aβ (Red) signal, and low temperature 4℃ (B). Colour yellow is the merged signal of EVs (Green) and Aβ (red). Scale bar= 20μm.

### 2.6. *In vitro* assessment of EV-Aβ association in response to other stress conditions and with proteins implicated in other neurodegenerative disorders

EVs are known to carry amyloid-β as an EV cargo and are often called in for causing the seeding effect in the progression of the AD pathology (15,33). Many studies have reported the amyloid beta aggregates to be topologically bound to the EVs surface membranes via glycolipid anchor. Transmission electron microscope (TEM) images have revealed that amyloid-β is attached to the surface of the EVs (34,35). To understand the interaction between amyloid-β and extracellular vesicles, we subjected the EVs and amyloid-beta peptides to different mechanical stresses like Ultrasonication and mild agitation. We also incubated EVs and amyloid-β together and assessed their association by TEM (Figure 6 A). In addition, confocal microscopy was used for further morphological characterization using Antibody-specific, precise and concise detection. EVs and amyloid-β did not show previously reported association when incubated without external mechanical stresses. However, when EVs and amyloid-β were ultrasonicated together, TEM images showed that amyloid-β was closely associated with EVs (Supplementary Figure 6A), and confocal images also showed a significant overlap between the two signals (Figure 6 B). This could imply that the ultrasonication altered the EV membrane affinity to Aβ and also caused Amyloid-β to aggregate (Supplementary Figure 6B). Furthermore, when we incubated EVs with amyloid-beta and then applied ultrasonication and mild agitation, we observed a distinct fibrillar structure similar to the BA structure in Figure 2A. However, this structure was not closely associated with the EVs. We also evaluated the fluorescence colocalization of two signals: AlexaFluor488 anti-CD9 and AlexaFluor647 anti-Amyloid-β. We measured Pearson’s correlation coefficient (PCC), which quantifies the overall association of two probes in an image (36). We observed a PCC value of 0.73 in Sonicated samples and a PCC of 0.54 in sonicated and agitated samples (Figure 6 C). These findings demonstrate an association between EVs and amyloid-β. However, whether the structural conformation of EVs, amyloid-β, or a combination of both drives this association remained uncertain.

**Figure 6:**
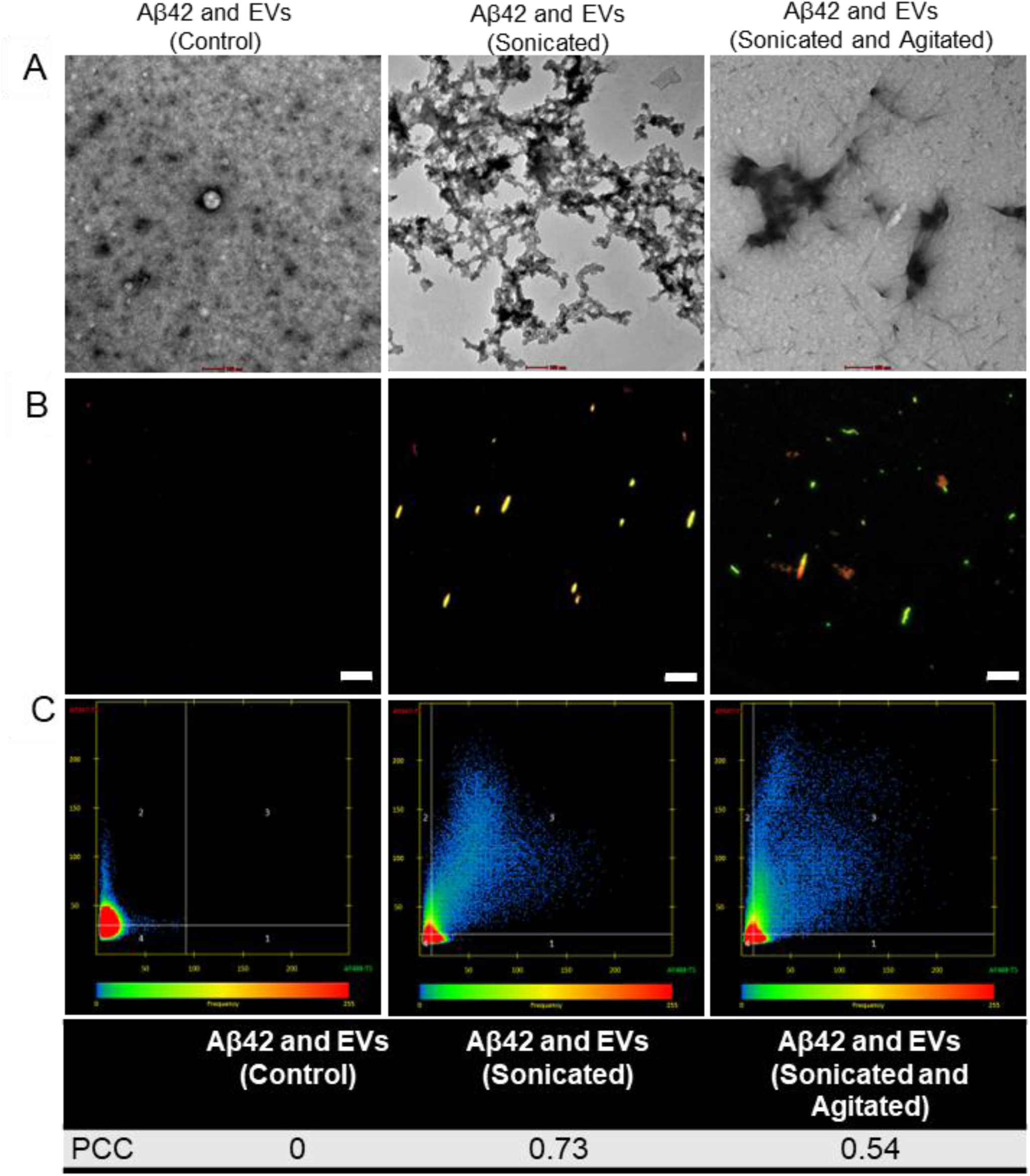
In-vitro assessment of amyloid-β and EV association. Imaging was performed at three key stages: Control (Without any stresses), Sonication only, and Sonication and agitation combined. TEM micrograph shows morphological characterization of different Aβ aggregated structures formed in the presence of EVs (A), CFM images for respective groups (B)and Colocalization analysis (C) showed maximum PCC in the post-sonication group (PCC= 0.73). Colour yellow is the merged signal of EVs (Green) and Aβ (red). Scale bar= 20μm.

Similarly, we conducted the experiment using alpha-synuclein, a hallmark protein associated with Parkinson’s disease. However, we did not observe any significant association with extracellular vesicles (EVs). (Supplementary figure 6C) This suggests that the observed selective interaction of altered EVs with amyloid-beta aggregates may be influenced by structural and size differences between the proteins.

### 2.7. Altered EV-Aβ coaggregates are internalized by cells and also localized at Amyloid Plaques

Confocal images revealed that EVs sequesters the amyloidβ aggregates, particularly the SA species. Consequently, co-incubation of BA with EVs did not reveal close proximation (Figure 7.1A, Supplementary Figure 7A). Similarly, the TEM micrograph shows the localization of amyloid-β aggregates (SA) around the EVs’ corona and when EVs +Aβ BA group, TEM images revealed the fibrils formed had clear filament twists (Figure 7.1 B). Notably, we concluded that the structural integrity of EVs also influenced the association with Aβ where more pronounced aggregates were formed with SA compared to BA (Supplementary figure 7B).

**Figure 7.1:**
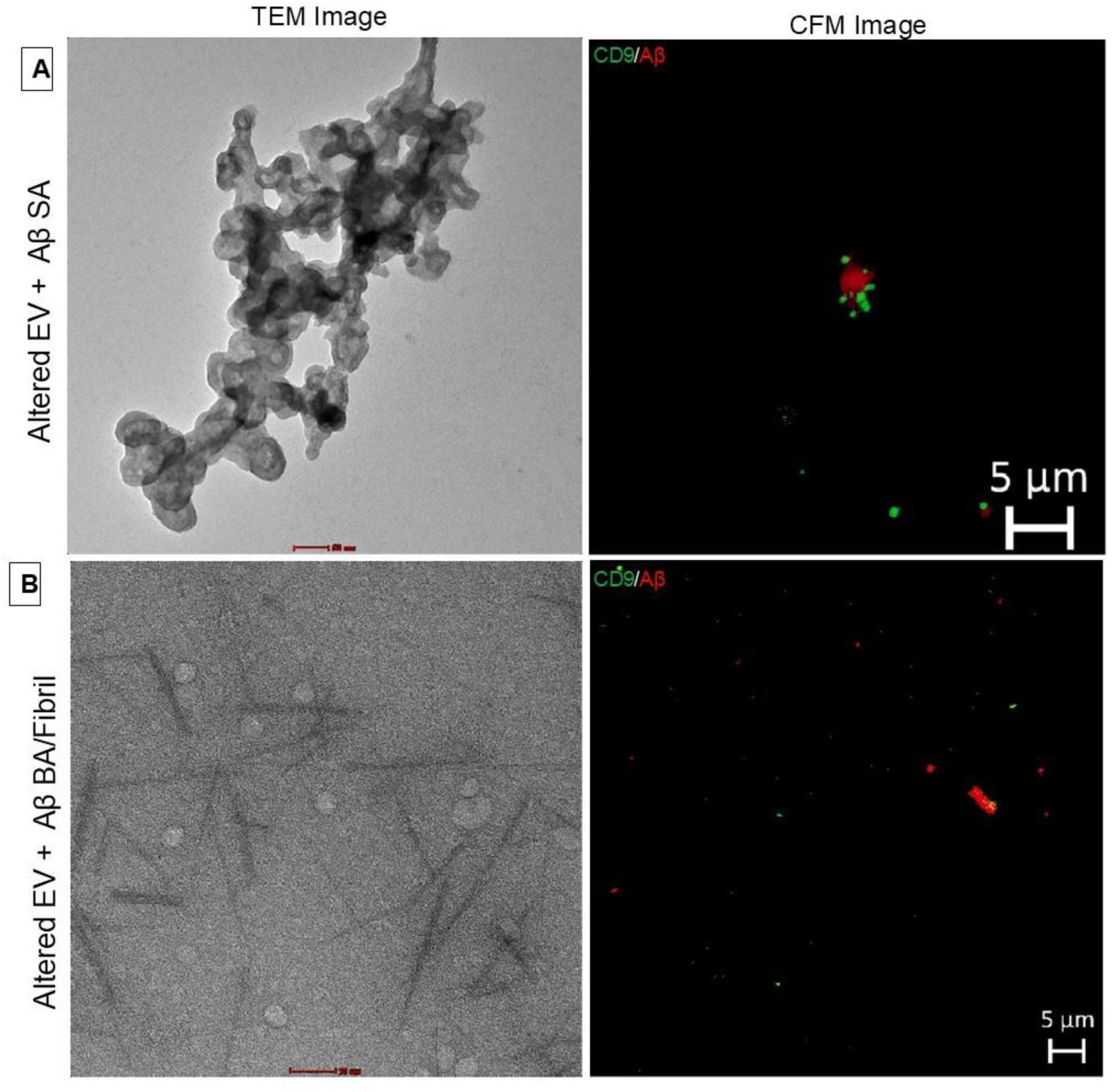
Comparison of interaction between EV and small amyloid-β aggregates (A) and big amyloid-β aggregates/Fibrils (B) at low temperature 4℃. The TEM, and CFM micrograph shows the EVs sequestering the SA group as opposed to no association between BA/Fibril (B).

**Figure 7.2.**
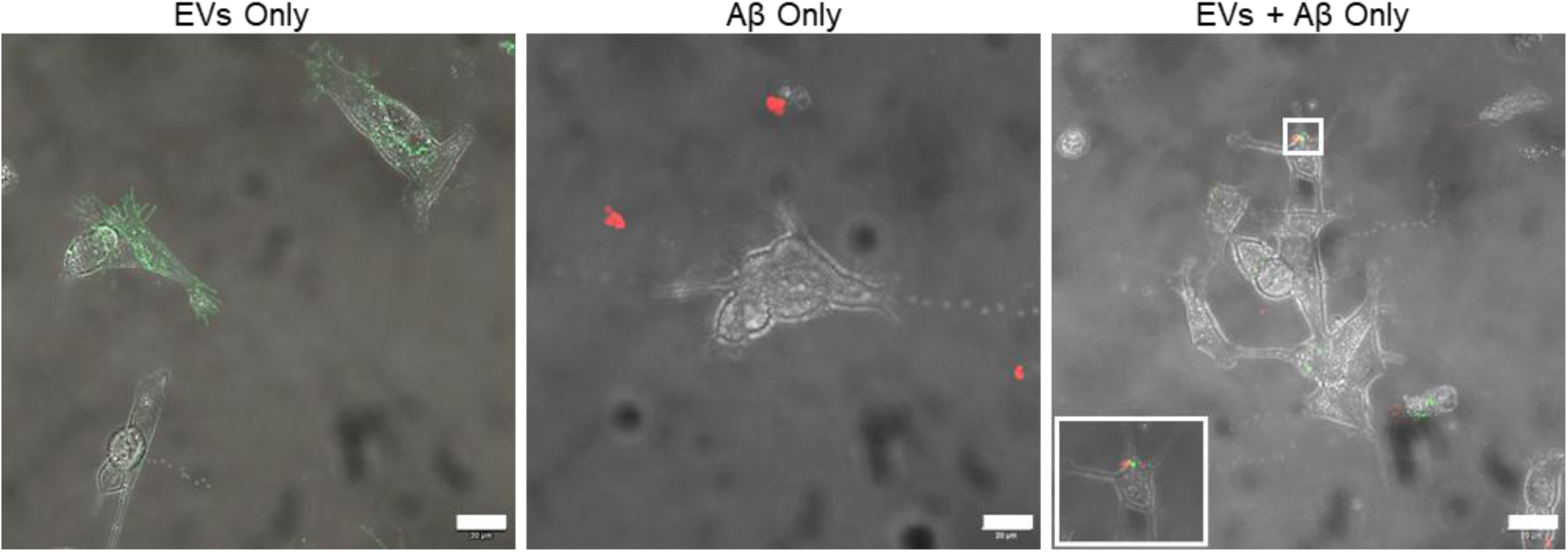
CFM image showing Alexa-Fluor-488 CD9 (Green) and Alexa-Fluor-647Amyloid-β (Red) signals of: (A) EVs only; (B) Aβ; and (C) EVs and Aβ together. Scale bar= 20μm.

**Figure 7.3:**
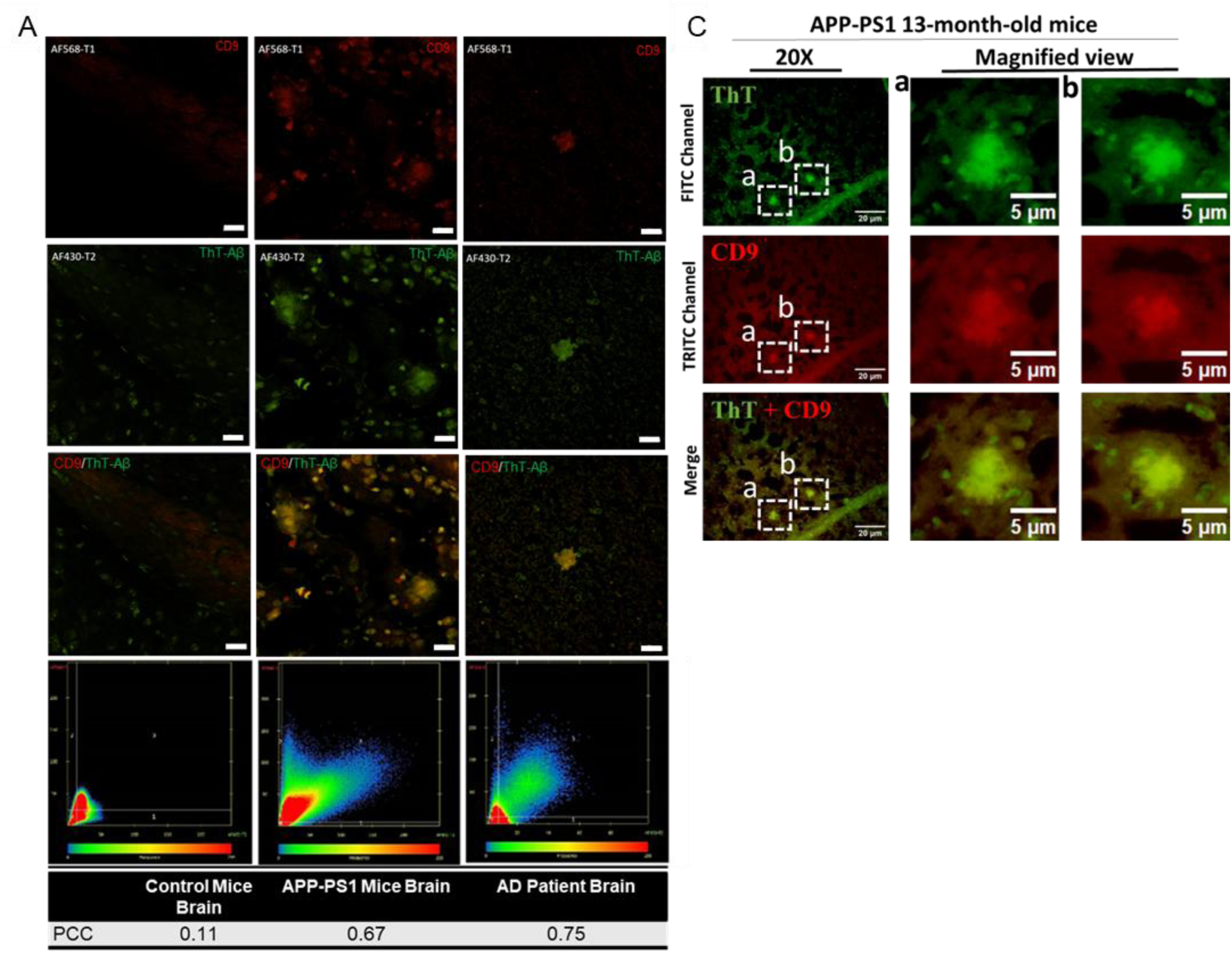
sEVs marker CD9 are enriched in amyloid-β plaque in both transgenic APP-PS1 mice and human brain section(A) and respective colocalization coefficient (B). In-vitro dual staining experimental images: (C) Control mice brain sections immunohistological staining with CD9 (antibody) and ThT stained Aβ plaques which are absent. APP mice brain section shows typical Amyloid-β plaques with discrete CD9 signals. Similarly, AD patient brain section also shows similar enrichment of CD9 signals around amyloid-β plaques. (n = 3 biologically independent samples) (Scale bar 20 μm; Zeiss Confocal microscope and NIS-Elements BR 4.30.00.64-bit fluorescence microscope). Images were captured at (For ThT; λex = 450 nm and λem = 490 nm. CD9-Rodamine TRITC-conjugated; λex = 550 nm and λem = 570 nm). Scale bar= 20μm.

To assess whether or not the altered EVs and Aβ aggregates are uptake by cells, EVs alone, amyloid-β aggregates alone, and altered EVs and amyloid-β coaggregates were labeled using Alexafluor488 anti-CD9 and Alexafluor647 anti-amyloid-β antibodies and were added to SH-SY5Y cells culture media. The cells were then extensively washed and examined by confocal scanning microscopy. Similar to EVs, altered EVs and amyloid-β coaggregates were found to be internalized by the cells. However, Aβ aggregates alone did not show internalization (Figure 7.2, Supplementary Figure 7C). Furthermore, the imaging of different optical sections of the cells revealed the presence of coaggregates in the plasma-membrane vicinity following their internalization. The internalization of only co-aggregates inside the cells could be attributed to EVs-mediated cellular uptake. Small red amyloid-β signals were detectable in confocal microscopy images outside the cells in amyloid-β alone, indicating that amyloid-β by itself does not enter cells. This suggests that only with the assistance of EVs can amyloid-β cross the cell membrane and reach the cell interior.

To determine whether these findings are relevant to AD pathology, we conducted immunohistochemistry using antibodies against the exosomal protein CD9 and performed ThT staining to label amyloid-β plaques in autopsy brain sections from Alzheimer’s disease patients, APP mice, and control mice (Figure 7.3A). The CD9 EV marker signal was observed surrounding small Aβ plaques, while the ThT signal was dispersed within the plaques in brain sections from both AD patients and APP-PS1 mice. In contrast, ThT staining was largely absent in the brain sections of control subjects. Pearson’s Correlation Coefficient for CD9-Aβ fluorescent labeled brain sections for: control mice were 0.11, APP-PS1 mice brain= 0.67, and for human AD brain= 0.75 suggesting enrichment of EVs around Amyloid-β plaques (Figure 7.3 B). Similarly, fluorescent images of the APP-PS1 mice brain reveal significant colocalization and accumulation of CD9 proteins in and around amyloid-β plaques (Figure 7.3 C). Extracellular deposition of Amyloid-β leads to the formation of senile plaques and characteristic exosomal CD-9 labeling observed throughout the vicinity of plaques provides a plausible mechanistic explanation that the altered EVs sequester small amyloid-β.

Congruent to the earlier findings reporting the enrichment of exosomal proteins around amyloid plaques in AD patients where it is involved in plaque formation, we further add that the altered sEVs preferentially sequester small amyloid-β aggregates and contribute to plaque nucleation and disease progression.

### 2.8. Assessment of EVs-Aβ co-aggregates in circulating plasma-derived EVs

Having determined that altered EVs preferentially sequester small amyloid-β aggregates, we investigated this association in circulating fluids, such as plasma, to assess whether these in vitro coaggregates can be detected. We isolated circulating EVs from Age-matched controls (AMC), and patients with mild cognitive impairment (MCI) and Alzheimer’s disease (AD), then incubated them with Anti-CD9 and Anti-Amyloidβ42 antibodies. The Confocal image showed no association between the two signals in circulating EVs, in contrast to our *in-vitro* findings (Figure 8A, Supplementary figure 8A). This is likely because extracellular amyloid-β aggregates cannot transverse the Blood-brain barrier (BBB) in association with circulating EVs. Next, we incubated EVs from Healthy controls with SA but observed no association (Figure 8B, Supplementary figure 8B). However, when AD EVs were co-incubated with SA, confocal images showed association between the two signals in AD patient samples corroborating with the *in-vitro* findings (Figure 8C, Supplementary figure 8C). This association may be due to the oxidative stress-induced conformational change in the EVs membrane making them more likely to associate with the Aβ SA aggregates.

**Figure 8:**
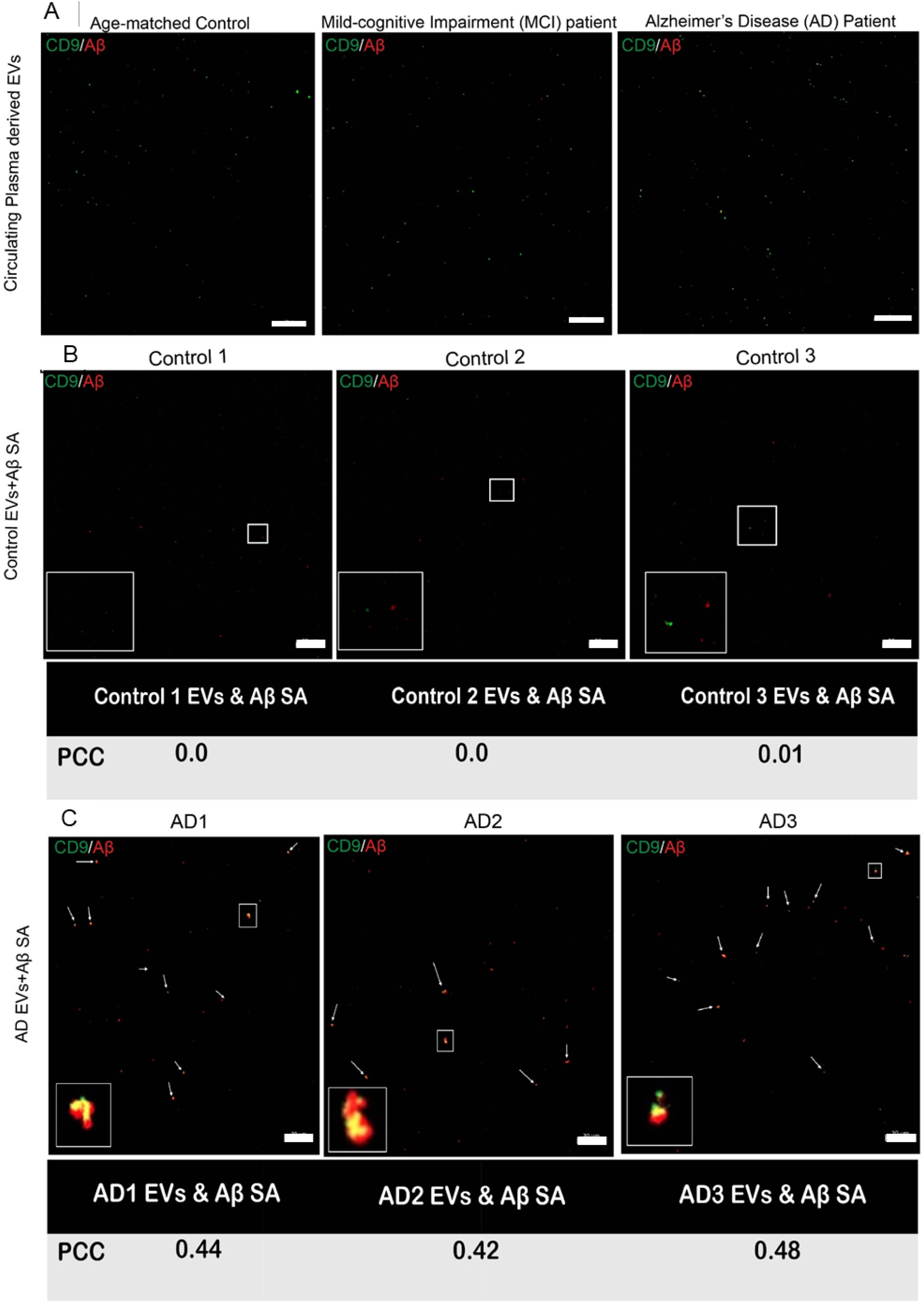
Circulating EVs derived from Control, MCI and AD patient does not show colocalization signal (A). Controls EVs incubated with small amyloid-β aggregates (B). AD EVs when incubated with small amyloid-β aggregates colocalize (White arrows) at low temperature 4℃ (C). Colour Yellow is the colocalized signal for EVs (Green)and Aβ (Red). Scale bar= 20μm.

**Figure 9:**
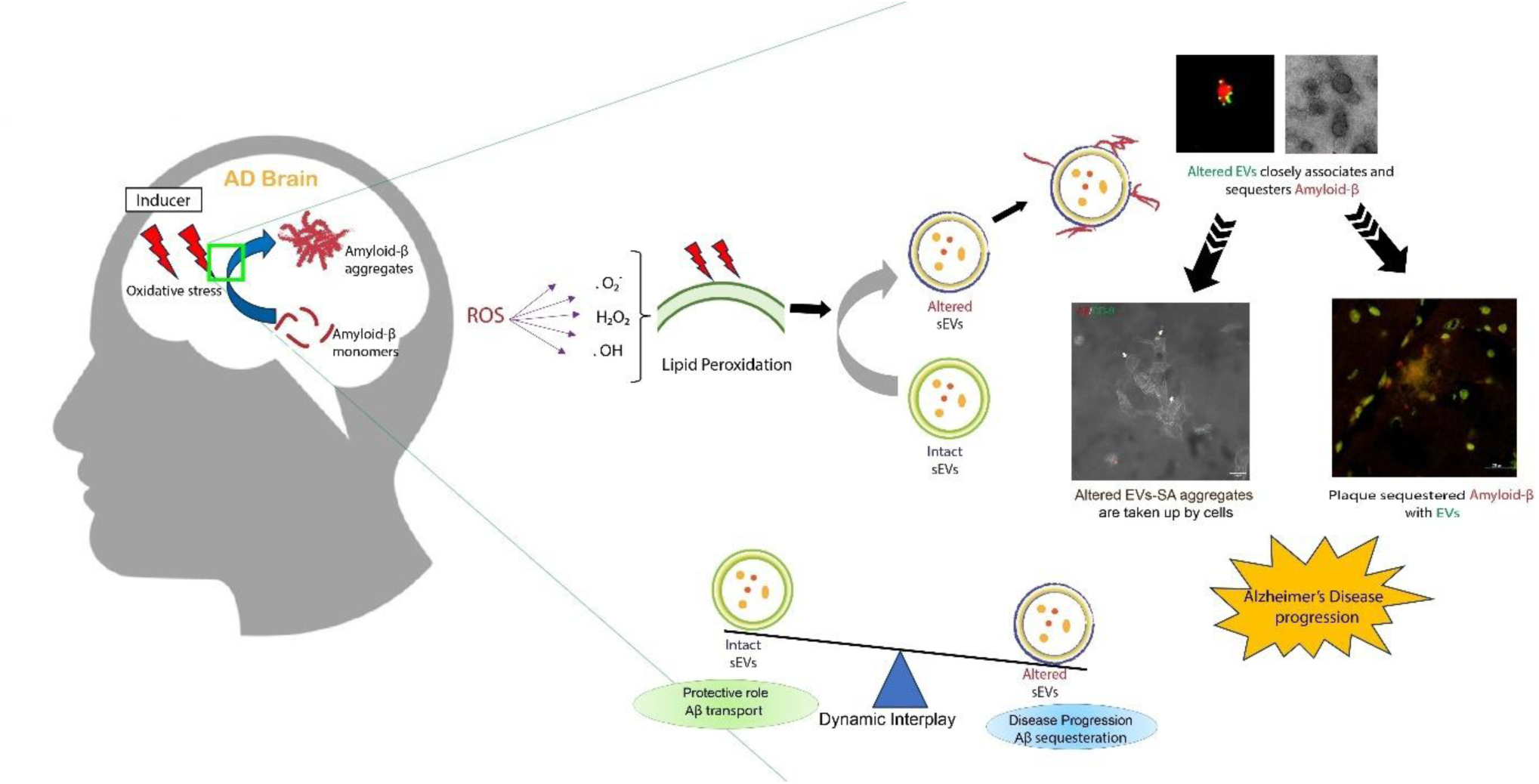
Mechanisms of Aβ Sequestration by stress-altered EVs and their implication in disease progression

## 3. Discussion

### 3.1. EVs Biological relevance

Cell-to-cell communication is mediated through various mechanisms, with extracellular vesicles playing a key role in the exchange of biomolecules secreted by cells. EVs are found in all biofluids and carry molecular signatures reflective of their cell of origin, which can influence various cellular processes under normal and pathological conditions (37). EVs typically have diameters ranging from 30 to 1000 nm and are categorized based on their size and biogenesis pathway into small extracellular vesicles (sEVs) (30–200 nm) or microvesicles (200–1000 nm) (38). EVs surface markers include Tetraspanin viz; CD63, CD81, CD9, and luminal markers, Tsg101, Alix. (39). In this study, EVs were isolated using a PEG-based chemical precipitation method followed by ultrafiltration, yielding analytical-grade EVs, consistent with the results observed in our previous studies (40–42). We characterized EVs by NTA to assess size distribution and particle concentration. Additionally, we used the immunoblot approach wherein EVs were analyzed for the expression of surface markers such as CD9, CD63 and CD81 and the presence of specific luminal markers like Tsg101.

### 3.2. Amyloid-β (Aβ) and EVs in Alzheimer’s Disease (AD)

Amyloid-β aggregates were prepared following the protocol by Lim et al. (2019) (35) to model species relevant to Alzheimer’s disease (AD) pathology by resuspending lyophilized HFIP-treated Aβ in 60mM NaOH, pH 7.4. Hexafluoroisopropanol (HFIP) treatment of amyloid-beta (Aβ) peptides is typically done to achieve a monomeric, non-aggregated state, which serves as a uniform starting point essential for controlled studies concerning amyloid-β peptides. Different-sized aggregates of Aβ-42 were generated, and their structural morphology and size distribution were confirmed through TEM, CFM, DLS, NTA, and Thioflavin-T (ThT) fluorescence. The EVs are known to carry amyloid-β (Aβ) as cargo and contribute to the seeding effect in Alzheimer’s disease (AD) progression (43). Previous studies have observed that Aβ aggregates can bind to EV membranes through electron microscopy (EM) (35). These studies have reported TEM-based examination of EV-Aβ interactions by immobilization on grids. There is a need to study these interactions in a physiological-like state, as we have observed alteration in EV membrane in response to external environmental conditions. Furthermore, the reported association between EVs and Aβ remains poorly understood. To investigate the interaction between EVs and Aβ, we employed an immune-fluorescence strategy followed by their visualization using confocal microscopy and TEM. TEM and confocal microscopy revealed that SA exhibited the highest binding affinity to EVs, with co-localization analysis yielding a PCC of 0.88 compared to minimal association with UA (PCC = 0.07) and BA (PCC = 0.02). This demonstrated that SA preferentially binds to EVs compared to other Aβ aggregates, and under normal conditions, EVs do not bind Aβ aggregates. Halipi *et al.* 2024 have extensively studied the intersection between EVs and Aβ (1–42) peptides, where authors have reported that EVs slow the aggregation of Aβ (1–42), primarily by inhibiting the fibril elongation phase (16). Similar conditions and concentrations were used in our work [EV concentrations of ∼10^9 particles/mL and Aβ (1–42) monomer concentration of 20 μM. We performed the temporal assessment of EVs-Aβ co-incubation for 24, 48, and 72 hours at both physiological temperatures, 37°C and 4°C to account for temperature-induced effects. EVs-Aβ co-incubation experiments at 37°C revealed a time-dependent increase in Aβ aggregate size, accompanied by EV aggregation confirming that physical conditions have a significant impact on the EVs membrane (31,32). This alteration of the EV membrane scaffold impacts the affinity to Aβ resulting in its sequestration whereas no major differences were observed in the experimental group at 4℃. Interestingly, subjecting EVs to mechanical stress such as ultrasonication before incubation at 4°C increased EVs association with Aβ. These findings indicate that alteration to the EVs membrane by mechanical (Ultrasonication and agitation) and physical (temperature) stress plays a critical role in Aβ sequestration.

### 3.3. Effect of Oxidative Stress on EVs and Aβ sequestration

We investigated whether oxidative stress, a well-recognized factor in Alzheimer’s disease (AD) pathogenesis, alters EVs through excessive production of reactive oxygen species (ROS), thereby influencing Aβ sequestration. To model this, we exposed EVs to a minimal dose of oxidative stress inducer i.e.,1 µM H_2_O_2_, for 24 hours and compared them to ultrasonicated EVs. Both treatments led to comparable structural distortions in EVs, as observed by transmission electron microscopy (TEM) and confocal microscopy, this was observed even at lower temperature conditions i.e., 4°C. These findings were consistent across temporally assessed groups-24, 48, and 72 hours whereby a gradual increase in the colocalization coefficient (PCC) indicating association of EV-Aβ signals was observed. This implies active sequestration by the altered EVs over time. We also observed that the structural morphology of EVs was distorted in both groups compared to control EVs. In addition to this, insight from lipidomic analysis of AD patients’ EVs demonstrated elevated levels of oxidized glycerophospholipid, which supports our inference that oxidative stress alters EV membrane scaffolds, influencing amyloid-β sequestration. Our results demonstrated that both oxidative stresses induced by H_2_O_2_ and mechanical (Ultrasonication) stress alter EVs membrane scaffold enhancing their affinity to Aβ which results in its sequestration.

### 3.4. Mechanisms of Aβ Sequestration by EVs and its role in disease progression

This study distinguishes between small aggregates (SA), which are preferentially sequestered by EVs, and larger aggregates (BA), which exhibit minimal interaction. This differentiation addresses the heterogeneity reported in TEM-based studies on EV-Aβ interactions. By studying these interactions in solution using an immunofluorescence approach and confocal microscopy, we avoided artifacts caused by immobilization on grids. Another study (13) have reported the prefibrillar Aβ aggregates binding preferentially with exosomes, however, the higher temperature conditions used in the study itself could be the driving force behind the EV-Aβ association due to EVs membrane alteration. Furthermore, enhanced binding of bigger Aβ42 aggregates (pre-fibrillar aggregates) to exosomes also does not hold good. This is due to the fact that structural flexibility and exposure of hydrophobic surfaces are critical factors that determine the capacity of oligomeric assemblies to cause cellular dysfunction, potentially resulting in outcomes such as neurodegeneration (44). A recent study reports that tau is tethered to the EV luminal membrane (45) as observed in the Cryo-EM imaging, inviting clarification behind this interaction as it leaves questions about the presence of tethering sites at EVs membrane, as tethering may require an adaptor or direct anchoring. Based on our observations, we propose that oxidative stress-induced conformational changes in EVs expose the limiting membrane, creating tethering sites for small aggregates. Studies have consistently shown that oligomers can interact with and permeabilize both synthetic lipid vesicles and cell membranes (46). Their high structural plasticity and hydrophobic surfaces enable oligomeric aggregates to penetrate cell membranes, traverse them, and reach the cell interior (47). Studies have shown that soluble oligomers from various types of amyloids increase lipid bilayer conductance, irrespective of their sequence. In contrast, fibrils and low molecular weight soluble species do not produce this effect. Notably, this increase in membrane conductance occurs without clear evidence of channel or pore formation or any ion selectivity (48). Untargeted lipidomic analysis revealed that lipid classes such as lysophosphatidylcholine (LPC), sphinganine (SPB), and ceramide (Cer) were highly dysregulated in stress-altered EVs compared to normal EVs, and lipid levels in the AD group closely mirrored those observed in EVs exposed to oxidative stress. Notably, studies show that Lysophosphatidylcholine (LPC) is elevated in oxidatively damaged tissues, and is also well-known for its role in the oligomer formation process of Aβ1-42 peptide, leading to a subsequent cascade of apoptosis and neuronal death (49–51). In our study, the small aggregates which represent oligomers, are preferentially sequestered by altered EVs enriched in these lipid classes such as LPC, SPB, which in turn promote EV-Aβ coaggregates that are subsequently internalized by cells, contributing to pathogenesis. This causes selective sequestration by altered EVs, highlighting their affinity for small aggregates (SA). Furthermore, SA may disrupt lipid membranes and internalize into neuronal cells, mediating toxicity. This process is likely to exacerbate neurodegeneration by triggering neuroinflammation and apoptotic pathway cascades thereby amplifying neuronal damage.

Furthermore, we also observed an enrichment of EVs CD9 marker around amyloid-β plaques where CD9 signals were localized around amyloid plaques in human AD and transgenic APP-PS1 mouse brains. This is consistent with previous findings showing the presence of exosomal proteins, such as flotillin-1 and Alix, near these plaques. Previous studies have reported that flotillin-1 labeled was most concentrated in areas containing senile plaques in the cerebral cortex, while the amyloid core of mature plaques was mostly negative for flotillin-1 (52) and an accumulation and enrichment of Alix were observed around amyloid plaques in AD patient’s brains (12). Our findings align with previous studies and demonstrate that altered EVs preferentially sequester small amyloid-β aggregates and contribute to plaque formation and disease progression.

We further examined the circulating EVs from age-matched controls (AMC), mild cognitive impairment (MCI), and Alzheimer’s disease (AD) for EVs and amyloid-β association. No association was observed likely because extracellular amyloid-β aggregates cannot transverse the blood-brain barrier (BBB). Similarly, EVs from healthy controls did not show an association when incubated with SA, in contrast, a significant association was observed when EVs from AD patients were incubated with SA, possibly due to a subset of altered EVs in circulation or disease-specific changes in EV properties.

### 3.5. Conclusions and Implications for Alzheimer’s Disease Progression

In our study, we observed that higher temperature conditions (31,32), H_2_O_2_-(53,54) and ultrasonication treatments led to significant distortion and self-aggregation of EVs. We observed structural changes in EVs subjected to these external stress exposures altered EV membrane scaffold that facilitated the sequestration of extracellular Aβ. Additionally, we validate the role of oxidative stress, driven by reactive oxygen species, as a key factor in Alzheimer’s disease (AD) pathogenesis, altering EVs membrane conformation and influencing Aβ aggregation. Notably, our study provides a clear demarcation of Aβ species— small aggregates (SA) preferentially sequestered by EVs. Our finding underscores the limitation of detecting extracellular Aβ aggregates in patient samples, likely due to blood-brain barrier (BBB) constraints. However, EVs from AD patients co-incubated with small Aβ aggregates (SA) exhibited a clear association, corroborating our in vitro results. This association likely stems from oxidative stress-induced alterations in EV membranes that enhance their affinity for Aβ SA. In summary, our study underscores the dual role of EVs in AD progression. While EVs typically support neuroprotection by clearing Aβ, altered EVs sequesters Aβ, promoting plaque formation, and crossing neuronal membranes to mediate toxicity, thereby driving disease progression.

This study provides a novel insight into the oxidative stress-mediated EV membrane conformational changes and their relevance in AD disease progression by sequestering more toxic SA and plaque nucleation. More research is necessary to comprehend this connection’s biological importance and investigate any potential implications for the onset of AD.

## 4. Materials & methods

### 4.1. Sample collection

Five millilitres of blood were collected from each participant via venipuncture into EDTA vials. Post-centrifugation of 2500 g for 20 minutes at 4 °C to pellet down the cells, plasma was collected, and subjected to centrifugation for 30 minutes at 4 °C and 10,000 g. Cleared plasma was aliquoted and stored at-80 °C. Post storage, the samples were centrifuged at 10,000 g after being thawed on ice and used for the downstream experiment. The study included healthy volunteers aged 25 to 30 with no family history of cancer, autoimmune diseases, neurodegenerative disorders, or amyloidosis, as well as Alzheimer’s disease patients diagnosed through neuropsychological assessments (ACE-III and MMSE) conducted by certified geriatric practitioners. Ethical approval for the study was granted under IECPG-213/20.04.2023, and all participants were enrolled after providing signed informed consent.

### 4.2. Isolation of sEVs

Small Extracellular vesicles (sEVs) were isolated using size exclusion chromatography (SEC) columns (Izon qEV1 70 nm Gen 2). We also isolated sEVs using the lab-developed filtration with precipitation method Bharti *et al* 2025 (In Press) (40–42). 1 ml of clarified plasma sample was used for isolating EVs from SEC columns. 1X PBS (ML116-500ML, HiMedia) was used to clean the column The fraction number 1-5 enriched in sEVs was subsequently used for downstream experiments.

### 4.3. Aβ42 aggregate formation

Aβ42 peptides (β-Amyloid (1–42) 1932-2-15) were purchased from DGpeptides Co., Ltd. 1 mg of peptide was dissolved in 1 mL of Hexafluoroisopropanol (HFIP) in a clean vial, then vortexed until fully dissolved. The resulting solution was aliquoted into 10 MCTs and sealed before flushing with nitrogen to evaporate the HFIP completely. After evaporation, the peptide was lyophilized before proceeding with any further processing. Different amyloid-β aggregates were prepared according to *Lim. et al. 2019* (35). Aβ42 peptides were resuspended in 60 mM NaOH and vortexed rigorously. To maintain the peptides at physiological pH, osmolarity, and ion concentration, the Aβ42 peptides were diluted to 20 µM, Un-aggregated Aβ42 (Aβ42 UA), pH 7.4 in 1X PBS Solution (Himedia, ML116). For making Aβ42 Small Aggregates (Aβ42 SA), 20 µM Aβ42 peptides, pH 7.4 were ultrasonicated using a benchtop bath sonicator at 50 kHz and 80 W for 5 minutes. For Aβ42 Big Aggregates (Aβ42 BA) preparation, after sonication step, it was incubated at 37°C with rapid shaking for 2 h. Aβ42 preparations were flash-frozen in liquid nitrogen and stored at −80°C.

### 4.4. α-synuclein aggregate formation

Recombinant human alpha-Synuclein protein (Met1–Ala140), expressed in *E. coli* (Cat# SP-485-500, R&D Systems), was used to induce aggregation following a modified protocol (55). A 20 μM working solution was prepared from a 60 μM stock in 10 mM Tris-Cl buffer (pH 7.4) containing 1 mM EDTA and 10 mM NaCl. The mixture was incubated at 37°C with rapid shaking in the presence of glass beads for seven days to promote fibril formation. Aggregation was confirmed using Transmission Electron Microscopy (TEM).

### 4.5. Thioflavin T (ThT) assay

Thioflavin T (ThT), Sigma was dissolved in phosphate-buffered saline (PBS) and filtered using a 0.2 µm syringe filter. ThT was then added to 20 µM Aβ42 UA, Aβ42 SA, and Aβ42 BA solutions, respectively, to achieve a final ThT concentration of 60 µM. The samples were incubated at 37°C for 2 hours. Following incubation, 40 µL of each ThT sample was transferred to a black 384-well opaque bottom plate. ThT fluorescence was measured at room temperature using a Spectramax i3x plate reader with an excitation wavelength of 450 nm and an emission range of 480 to 600 nm.

### 4.6. Western blot

All samples were normalized according to the initial volume of biofluid input, i.e. 180µl (41,42). Then, 20 μL was loaded to run on an 8–12% SDS PAGE after the sEVs sample and the sample loading dye (2 × Laemmle Sample buffer) were combined. Following SDS-PAGE, the protein from the gel was wet transferred onto a 0.22 μm PVDF membrane (1,620,177, BioRad). Membrane was blocked with 3% bovine serum albumin (BSA) (D0024, BioBasic) in Tris (TB0194, BioBasic) base saline containing 0.1% of Tween 20 (65,296, SRL Chem) (TBST). Blot was incubated with primary antibodies against CD63 (10628D, Invitrogen), CD81 (PA5-86,534, Invitrogen), TSG101 (MA1-23,296, Invitrogen) and CD9 (PA5-86534, Invitrogen) were incubated at 4 °C for the entire night. Before being incubated at room temperature with HRP-conjugated secondary antibodies, anti-rabbit (AB6721, Abcam), and anti-mouse (31,430, Invitrogen), the membranes were washed three times with TBST buffer. Apolipoprotein levels were also assessed to check the level of protein co-isolates.The blot was developed using the Femto LUCENT™ PLUS-HRP kit (AD0023, GBiosciences) to visualize the protein bands through enhanced chemiluminescence.

### 4.7. Nanoparticle tracking analysis (NTA)

The NTA of sEVs was performed after 5000-fold dilution in 1X-PBS buffer. One milliliter of a diluted sEVs sample was added to the sample chamber of the ZetaView Twin system (Particle Metrix, Germany). The following settings were applied during three cycles of scanning eleven different cell locations: high video setting, autofocus: focus, shutter: 150, camera sensitivity: 80, and cell temperature 25 °C. A total of sixty frames were gathered for each position. The built-in ZetaView Software 8.05.12 (Particle Metrix, Germany) was used for analysis, with CMOS cameras for recording. The least particle size was 10 nm, the maximum particle size was 1000 nm, and the minimum particle brightness was 30.

### 4.8. Dynamic Light Scattering (DLS)

1µl of Aβ42 preparations (Aβ42 UA, Aβ42 SA, Aβ42 BA) was added to 1000µl of 1XPBS, vortexed, and transferred to a precleaned sample cuvette. Hydrodynamic size (Rh) distribution measurements were performed by a zeta particle size analyzer (Malvern).

### 4.9. Co-incubation of sEVs with different Aβ42 preparations

#### 4.9.1. Assessment of EVs-amyloid beta association

Approximately 9.0×10^8^ particles of sEVs were introduced to 20 µM Aβ42 UA solution. The mixture was subjected to sonication using a benchtop sonicator at 80 W power and 60 kHz frequency. Following sonication, the samples were incubated for 2 hours with agitation.

#### 4.9.2. Evaluation of sEVs’ affinity with different Amyloid-beta aggregates

Aβ42 UA, Aβ42 SA, and Aβ42 BA, each at a concentration of 20 µM, were individually combined with approximately 6.0×10^8^ particles of sEVs. The mixtures were incubated for 2 hours at 25°C. Imaging was performed following the incubation period.

#### 4.9.3. Assessment of Amyloid beta-sEVs association in a time-dependent manner

Aβ42 UA was incubated with ∼10^9^ EV particles at room temperature and 4°C. The association between Aβ42 UA and sEVs was assessed by imaging at different time points, specifically after 24, 48, and 72 hours of incubation.

#### 4.9.4. Amyloid Beta Association with Ultrasonicated sEVs

A 20 µM solution of Aβ42 SA was added to approximately 9.0×10^8^ EV particles pretreated by ultrasonication using a benchtop sonicator. The combined sample was then incubated at 4°C, and imaging was performed at 24, 48, and 72-hour time points to assess the interaction between Aβ42 SA and the sonication-induced disrupted sEVs.

#### 4.9.5. Amyloid Beta Association with sEVs in the presence of H_2_O_2_

∼9.0×10^8^ EV particles were combined with 20 µM Aβ42 SA in the presence of 100 µM hydrogen peroxide (H₂O₂). The mixture was incubated at 4°C and imaged at two-time points: immediately (0 hours) and after 24 hours.

### 4.10. Transmission electron microscopy

The ultrastructural morphology of sEVs was studied using transmission electron microscopy. The isolated sEVs were diluted (1:1000) with filtered 1× PBS. Similarly, for the characterization of different Aβ42 peptides and aggregates, 20 µM solution of Aβ42 UA, Aβ42 SA, and Aβ42 BA were diluted to 1000-fold in filtered 1× PBS. Finally, sEVs and Aβ42 preparations were co-incubated and the samples were diluted 2500X in 1X PBS. Subsequently, small drops (50µl) of the diluted sample suspension were adsorbed at room temperature for 15 minutes using a 300-mesh carbon-coated copper grid (01843, Ted Pella). Following this, grids were gently dabbed to remove excess liquid and then positioned onto a 20 µL droplet of filtered MilliQ water. As a negative stain, 2% aqueous uranyl acetate solution (81,405, SRL Chem) was used for 10 seconds. The grids were blot-dried and examined with a Thermo Scientific Talos S transmission electron microscope.

### 4.11. Confocal microscopy

#### 4.11.1. Characterization of sEVs

20µl of isolated sEVs were incubated for 2 hours at 25⁰C with 0.4µl of Human CD9 Alexa Fluor® 488-conjugated antibody (R&D systems, FAB1880G) and imaged using a Zeiss LSM980 confocal microscope (Carl Zeiss Microscopy).

#### 4.11.2. sEVs and Aβ42 preparations co-incubated

sEVs and Aβ42 suspension was labeled using Human CD9 Alexa Fluor® 488-conjugated Antibody (R&D systems, FAB1880G) for sEVs and Alexa Fluor® 647 Anti-beta Amyloid 1-42 antibody [mOC64] (Abcam, ab300742) in final concentration of 2%(v/v) and 0.5%(v/v) of the total suspension respectively. sEVs and the aggregate suspension were incubated for 2 hours with both antibodies at 25⁰C before mounting on labolene-cleaned glass slides (Blue Star) and covered with 18mm;10Gms glass covers (Blue Star), and were imaged on Zeiss LSM980 system using the 40X objective. The acquisition parameters were kept constant for the same set of experiments.

### 4.12. dSTORM Microscopy

STORM images were acquired using a custom-built setup with a Nikon Ti2E microscope body. Briefly, Aβ42 labelled with Alexa Fluor® 647 Anti-beta Amyloid 1-42 antibody [mOC64] (Abcam, ab300742) was excited with a 647 laser (MPB Communications). Upon excitation, the emitted light was collected by Nikon 100x oil immersion TIRF-SR objective (NA 1.49) and a laser quad band set with emission filter (TRF89902-EMET-405/488/561/647 nm laser Quad Band Set for TIRF application; Chroma). An EM-CCD camera was used for capturing images and hence, image localization, at an exposure time of 20 milliseconds per frame for 40,000 frames for each STORM image. Reconstruction of the image was done using custom-written software (Insight3, provided by B. Huang, University of California, San Francisco, CA, USA) (56).

To understand the degree of interaction with SEC-isolated EVs labeled using Human CD9 Alexa Fluor® 488-conjugated Antibody, Total Illumination Reflection Microscopy was performed on the same sample, using the same microscope. The TIRF images were captured with the EM-CCD camera at an exposure time of 100 milliseconds per frame for 10 frames. Upon acquisition, the TIRF images were overlaid on the STORM images with the image coordinates in place to study the interaction between the aggregate and the extracellular vesicles.

### 4.13. Lipidomic study

Lipids were isolated using modified Folch method which employed a biphasic solvent system composed of chloroform and methanol in a 2:1(v/v) ratio (57). The LC-MS analysis was performed on a high-resolution Orbitrap Exploris 120 mass spectrometer (Thermo Fisher Scientific) equipped with a Vanquish UHPLC (Thermo Fisher Scientific). The acquired data was processed using Compound Discover software (3.2.1 version; Thermo Fisher Scientific), and metabolite screening was done as an untargeted approach.

### 4.14. Cell-internalization of EV-Aβ coaggregates

SH-SY5Y human neuroblastoma cells were grown in DMEM/F-12 (Dulbecco’s Modified Eagle Medium/Nutrient Mixture F-12), Gibco™ (Cat: 11320033), supplemented with 10% qualified fetal bovine serum (FBS), Gibco™ (Cat: 10270106), and 1% Antibiotic-Antimycotic (100X), Cat: 15240062, under standard conditions at 37°C in a humidified atmosphere containing 5% CO₂. Cells seeded on Nunc™ Glass Bottom Dishes Cat: 150680 and were first treated for 60 min at 37 °C with EVs alone, Aβ aggregates alone, and EV-Aβ coaggregates. A total of 3µg protein was used. EVs and Aβ were labeled with primary antibodies conjugated with fluorophore namely, AlexaFluor647 anti-amyloidβ and AlexaFluor488 anti-CD9 antibodies. Spent media was replaced with fresh media before imaging.

### 4.15. Brain tissue staining protocol

Brain specimens were post-fixed in 4% paraformaldehyde for 72 hours, then transferred to 15% sucrose in PBS (Cat # S5-3, Fisher Scientific) until tissue settled to the bottom. Subsequently, tissues were moved to 30% sucrose in PBS until they sank to the bottom, indicating complete saturation. The tissues were embedded in an OCT compound and sectioned at a thickness of 12–15 µm using a cryotome. Cryo-sectioned brain slices were obtained from 13-month-old APPswe-PSEN1 mice (male) and age-matched wild-type (WT) mice (male). Brain slices were deparaffinized in xylene and rehydrated through sequential washes in 90%, 75%, 50%, and 30% ethanol, followed by a final rinse in 1× PBS, with each step lasting 5 minutes. EVs were labeled using a primary CD9 antibody (Catalog # PA5-86534, Invitrogen), followed by a TRITC-conjugated secondary antibody (Goat anti-Rabbit IgG (H+L), Catalog # A16101). Amyloid-beta (Aβ) deposits in the brain tissue sections were stained with 0.1% aqueous ThT solution. The sections were incubated for 60 minutes and then washed three times with PBS, 5 minutes per wash. Prepared tissue slices were mounted and visualized using a laser confocal microscope (LSM980, Carl Zeiss) at 40× magnification.

### 4.16. Colocalization and Statistical analysis

All confocal images were processed using the Zen 3.9 image processing software. Small extracellular vesicles (sEVs) labeled with Human CD9 Alexa Fluor® 488-conjugated antibody were visualized in the green channel. In contrast, amyloid-β aggregates, labeled with recombinant Alexa Fluor® 647 anti-beta amyloid 1-42 antibody, were observed in the red channel. Colocalized signals appeared yellow. In addition to visual colocalization, pixel-by-pixel colocalization analysis was performed using Zen 3.9. The intensity of the one-color channel was plotted against the intensity of the second color for each pixel in a scatterplot to graphically represent colocalization. Background noise was removed by applying threshold values determined by the Costes method (58). Colocalization was quantitatively assessed using Pearson’s Correlation Coefficient (PCC), ranging from +1 (indicating perfectly linearly related intensities) to-1 (indicating perfectly inversely related intensities), and was also used to quantify the strength of the correlation between the fluorescence intensities of the two channels (36,59).

## Acknowledgement

This work was supported by the Indian Council of Medical Research (ICMR: funding number 2020–1194) and the Department of Health Research (DHR: GIA/2020/000595). S.R. gratefully acknowledges the CSIR (09/0006(0519)/2019-EMR-I) for providing the fellowship. We thank Dr. Sarika Gupta, National Institute of Immunology, for providing the APP mice brain tissue samples for histological staining. We extend our gratitude to Dr. Anita Mahadevan, Coordinator of the Human Brain Bank (NIMHANS), for providing autopsy brain tissue samples from clinically diagnosed Alzheimer’s disease patients for histological staining. We are very thankful to Rishabh Singh and Umar Bin Tabrez, for their valuable assistance.

## Authors’ contributions

SK conceptualised and designed the study. SZ, SR, and SDC performed major experiments and data analysis. SR, SZ, and KR wrote the manuscript. HR and GPM participated in animal experiments. SB and NM acquired and analysed SRM data. AG and PC participated in AD patient recruitment and clinical assessment. TJR, NR, and FN performed initial proof. All authors participated in the finalisation of the manuscript.

## Data Availability Statement

All data are available in the main text or the supplementary materials.

## Ethical clearance

Ethical approval for the study was granted under IECPG-213/20.04.2023, and all participants were enrolled after providing signed informed consent.

## References

1. Blennow K, Leon MJ de, Zetterberg H. Alzheimer’s disease. The Lancet. 2006 Jul 29;368(9533):387–403.

2. Selkoe DJ. Alzheimer’s disease. Cold Spring Harb Perspect Biol. 2011 Jul 1;3(7):a004457.

3. Gu L, Guo Z. Alzheimer’s Aβ42 and Aβ40 peptides form interlaced amyloid fibrils. J Neurochem. 2013 Aug;126(3):305–11.

4. Kalluri R, LeBleu VS. The biology, function, and biomedical applications of exosomes. Science. 2020 Feb 7;367(6478):eaau6977.

5. van Niel G, D’Angelo G, Raposo G. Shedding light on the cell biology of extracellular vesicles. Nat Rev Mol Cell Biol. 2018 Apr;19(4):213–28.

6. Rastogi S, Sharma V, Bharti PS, Rani K, Modi GP, Nikolajeff F, et al. The Evolving Landscape of Exosomes in Neurodegenerative Diseases: Exosomes Characteristics and a Promising Role in Early Diagnosis. Int J Mol Sci. 2021 Jan 4;22(1):440.

7. Wang Y, Balaji V, Kaniyappan S, Krüger L, Irsen S, Tepper K, et al. The release and trans-synaptic transmission of Tau via exosomes. Molecular Neurodegeneration. 2017 Jan 13;12(1):5.

8. Laulagnier K, Javalet C, Hemming FJ, Chivet M, Lachenal G, Blot B, et al. Amyloid precursor protein products concentrate in a subset of exosomes specifically endocytosed by neurons. Cell Mol Life Sci. 2018 Feb 1;75(4):757–73.

9. Yuyama K, Sun H, Sakai S, Mitsutake S, Okada M, Tahara H, et al. Decreased Amyloid-β Pathologies by Intracerebral Loading of Glycosphingolipid-enriched Exosomes in Alzheimer Model Mice *. Journal of Biological Chemistry. 2014 Aug 1;289(35):24488–98.

10. Willén K, Edgar JR, Hasegawa T, Tanaka N, Futter CE, Gouras GK. Aβ accumulation causes MVB enlargement and is modelled by dominant negative VPS4A. Molecular Neurodegeneration. 2017 Aug 23;12(1):61.

11. Lee JH, Yang DS, Goulbourne CN, Im E, Stavrides P, Pensalfini A, et al. Faulty autolysosome acidification in Alzheimer’s disease mouse models induces autophagic build-up of Aβ in neurons, yielding senile plaques. Nat Neurosci. 2022 Jun;25(6):688–701.

12. Rajendran L, Honsho M, Zahn TR, Keller P, Geiger KD, Verkade P, et al. Alzheimer’s disease β-amyloid peptides are released in association with exosomes. Proceedings of the National Academy of Sciences of the United States of America. 2006 Jul 12;103(30):11172.

13. Lim CZJ, Zhang Y, Chen Y, Zhao H, Stephenson MC, Ho NRY, et al. Subtyping of circulating exosome-bound amyloid β reflects brain plaque deposition. Nat Commun. 2019 Mar 8;10(1):1144.

14. Yuyama K, Sun H, Mitsutake S, Igarashi Y. Sphingolipid-modulated Exosome Secretion Promotes Clearance of Amyloid-β by Microglia. Journal of Biological Chemistry. 2012 Mar;287(14):10977– 89.

15. Yuyama K, Sun H, Usuki S, Sakai S, Hanamatsu H, Mioka T, et al. A potential function for neuronal exosomes: Sequestering intracerebral amyloid-β peptide. FEBS Letters. 2015;589(1):84–8.

16. Halipi V, Sasanian N, Feng J, Hu J, Lubart Q, Bernson D, et al. Extracellular Vesicles Slow Down Aβ(1–42) Aggregation by Interfering with the Amyloid Fibril Elongation Step. ACS Chemical Neuroscience [Internet]. 2024 Feb 26 [cited 2024 Sep 10]; Available from: https://pubs.acs.org/doi/full/10.1021/acschemneuro.3c00655

17. Markoutsa E, Mayilsamy K, Gulick D, Mohapatra SS, Mohapatra S. Extracellular vesicles derived from inflammatory-educated stem cells reverse brain inflammation—implication of miRNAs. Molecular Therapy. 2022 Feb 2;30(2):816–30.

18. Sobue A, Komine O, Yamanaka K. Neuroinflammation in Alzheimer’s disease: microglial signature and their relevance to disease. Inflammation and Regeneration. 2023 May 10;43(1):26.

19. Guo T, Zhang D, Zeng Y, Huang TY, Xu H, Zhao Y. Molecular and cellular mechanisms underlying the pathogenesis of Alzheimer’s disease. Molecular Neurodegeneration. 2020 Jul 16;15(1):40.

20. Dhapola R, Beura SK, Sharma P, Singh SK, HariKrishnaReddy D. Oxidative stress in Alzheimer’s disease: current knowledge of signaling pathways and therapeutics. Mol Biol Rep. 2024 Jan 2;51(1):48.

21. Cummings JL, Osse AML, Kinney JW. Alzheimer’s Disease: Novel Targets and Investigational Drugs for Disease Modification. Drugs. 2023 Oct 1;83(15):1387–408.

22. Singh G, Kumar S, Panda SR, Kumar P, Rai S, Verma H, et al. Design, Synthesis, and Biological Evaluation of Ferulic Acid-Piperazine Derivatives Targeting Pathological Hallmarks of Alzheimer’s Disease. ACS Chem Neurosci. 2024 Aug 7;15(15):2756–78.

23. Shankar G, Praveen Kumar C, Yadav M, Ghosh A, Panda SR, Banerjee A, et al. Discovery of novel substituted (Z)-N′-hydroxy-3-(3-phenylureido)benzimidamide derivatives as multifunctional molecules targeting pathological hallmarks of Alzheimer’s disease. European Journal of Medicinal Chemistry. 2024 Dec 15;280:116959.

24. Duan J, Huang Z, Qin S, Li B, Zhang Z, Liu R, et al. Oxidative stress induces extracellular vesicle release by upregulation of HEXB to facilitate tumour growth in experimental hepatocellular carcinoma. J Extracell Vesicles. 2024 Jul;13(7):e12468.

25. Chiaradia E, Tancini B, Emiliani C, Delo F, Pellegrino RM, Tognoloni A, et al. Extracellular Vesicles under Oxidative Stress Conditions: Biological Properties and Physiological Roles. Cells. 2021 Jul 12;10(7):1763.

26. Taniguchi A, Sasaki D, Shiohara A, Iwatsubo T, Tomita T, Sohma Y, et al. Attenuation of the aggregation and neurotoxicity of amyloid-β peptides by catalytic photooxygenation. Angew Chem Int Ed Engl. 2014 Jan 27;53(5):1382–5.

27. Subramaniam R, Roediger F, Jordan B, Mattson MP, Keller JN, Waeg G, et al. The lipid peroxidation product, 4-hydroxy-2-trans-nonenal, alters the conformation of cortical synaptosomal membrane proteins. J Neurochem. 1997 Sep;69(3):1161–9.

28. Castegna A, Lauderback CM, Mohmmad-Abdul H, Butterfield DA. Modulation of phospholipid asymmetry in synaptosomal membranes by the lipid peroxidation products, 4-hydroxynonenal and acrolein: implications for Alzheimer’s disease. Brain Research. 2004 Apr;1004(1–2):193–7.

29. Michalska P, León R. When It Comes to an End: Oxidative Stress Crosstalk with Protein Aggregation and Neuroinflammation Induce Neurodegeneration. Antioxidants (Basel). 2020 Aug 12;9(8):740.

30. Hallal S, Tűzesi Á, Grau GE, Buckland ME, Alexander KL. Understanding the extracellular vesicle surface for clinical molecular biology. J Extracell Vesicles. 2022 Oct;11(10):e12260.

31. Görgens A, Corso G, Hagey DW, Jawad Wiklander R, Gustafsson MO, Felldin U, et al. Identification of storage conditions stabilizing extracellular vesicles preparations. Journal of Extracellular Vesicles. 2022;11(6):e12238.

32. Sivanantham A, Jin Y. Impact of Storage Conditions on EV Integrity/Surface Markers and Cargos. Life. 2022 May;12(5):697.

33. Accelerated release of exosome-associated GM1 ganglioside (GM1) by endocytic pathway abnormality: another putative pathway for GM1-induced amyloid fibril formation. [cited 2024 Aug 22]; Available from: https://onlinelibrary.wiley.com/doi/10.1111/j.1471-4159.2007.05128.x

34. Rajendran L, Honsho M, Zahn TR, Keller P, Geiger KD, Verkade P, et al. Alzheimer’s disease beta-amyloid peptides are released in association with exosomes. Proc Natl Acad Sci U S A. 2006 Jul 25;103(30):11172–7.

35. Lim CZJ, Zhang Y, Chen Y, Zhao H, Stephenson MC, Ho NRY, et al. Subtyping of circulating exosome-bound amyloid β reflects brain plaque deposition. Nat Commun. 2019 Mar 8;10(1):1144.

36. Dunn KW, Kamocka MM, McDonald JH. A practical guide to evaluating colocalization in biological microscopy. Am J Physiol Cell Physiol. 2011 Apr;300(4):C723–42.

37. Bellingham SA, Guo BB, Coleman BM, Hill AF. Exosomes: Vehicles for the Transfer of Toxic Proteins Associated with Neurodegenerative Diseases? Front Physio [Internet]. 2012 [cited 2024 Feb 9];3. Available from: http://journal.frontiersin.org/article/10.3389/fphys.2012.00124/abstract

38. Akers JC, Gonda D, Kim R, Carter BS, Chen CC. Biogenesis of extracellular vesicles (EV): Exosomes, microvesicles, retrovirus-like vesicles, and apoptotic bodies. Journal of neuro-oncology. 2013 May;113(1):1–11.

39. Mathew B, Mansuri MS, Williams KR, Nairn AC. Exosomes as Emerging Biomarker Tools in Neurodegenerative and Neuropsychiatric Disorders—A Proteomics Perspective. Brain Sci. 2021 Feb 19;11(2):258.

40. Rastogi S, Rani K, Rai S, Singh R, Bharti PS, Sharma V, et al. Fluorescence-tagged salivary small extracellular vesicles as a nanotool in early diagnosis of Parkinson’s disease. BMC Medicine. 2023 Sep 4;21(1):335.

41. Rai S, Bharti PS, Singh R, Rastogi S, Rani K, Sharma V, et al. Circulating plasma miR-23b-3p as a biomarker target for idiopathic Parkinson’s disease: comparison with small extracellular vesicle miRNA. Front Neurosci [Internet]. 2023 Nov 15 [cited 2024 Aug 22];17. Available from: https://www.frontiersin.org/journals/neuroscience/articles/10.3389/fnins.2023.1174951/full

42. Singh R, Rai S, Bharti PS, Zehra S, Gorai PK, Modi GP, et al. Circulating small extracellular vesicles in Alzheimer’s disease: a case–control study of neuro-inflammation and synaptic dysfunction. BMC Medicine. 2024 Jun 20;22(1):254.

43. Yuyama K, Sun H, Usuki S, Sakai S, Hanamatsu H, Mioka T, et al. A potential function for neuronal exosomes: sequestering intracerebral amyloid-β peptide. FEBS Lett. 2015 Jan 2;589(1):84–8.

44. Benilova I, Karran E, De Strooper B. The toxic Aβ oligomer and Alzheimer’s disease: an emperor in need of clothes. Nat Neurosci. 2012 Mar;15(3):349–57.

45. Fowler SL, Behr TS, Turkes E, O’Brien DP, Cauhy PM, Rawlinson I, et al. Tau filaments are tethered within brain extracellular vesicles in Alzheimer’s disease. Nat Neurosci. 2024 Nov 21;1–9.

46. Bucciantini M, Calloni G, Chiti F, Formigli L, Nosi D, Dobson CM, et al. Prefibrillar amyloid protein aggregates share common features of cytotoxicity. J Biol Chem. 2004 Jul 23;279(30):31374–82.

47. Campioni S, Mannini B, Zampagni M, Pensalfini A, Parrini C, Evangelisti E, et al. A causative link between the structure of aberrant protein oligomers and their toxicity. Nat Chem Biol. 2010 Feb;6(2):140–7.

48. Kayed R, Sokolov Y, Edmonds B, McIntire TM, Milton SC, Hall JE, et al. Permeabilization of Lipid Bilayers Is a Common Conformation-dependent Activity of Soluble Amyloid Oligomers in Protein Misfolding Diseases. Journal of Biological Chemistry. 2004 Nov;279(45):46363–6.

49. Prabutzki P, Schiller J, Engel KM. Phospholipid-derived lysophospholipids in (patho)physiology. Atherosclerosis. 2024 Nov 1;398:118569.

50. Sheikh AM, Michikawa M, Kim SU, Nagai A. Lysophosphatidylcholine increases the neurotoxicity of Alzheimer’s amyloid β1-42 peptide: role of oligomer formation. Neuroscience. 2015 Apr 30;292:159–69.

51. Law SH, Chan ML, Marathe GK, Parveen F, Chen CH, Ke LY. An Updated Review of Lysophosphatidylcholine Metabolism in Human Diseases. Int J Mol Sci. 2019 Mar 6;20(5):1149.

52. Kokubo H, Lemere CA, Yamaguchi H. Localization of flotillins in human brain and their accumulation with the progression of Alzheimer’s disease pathology. Neuroscience Letters. 2000 Aug 25;290(2):93–6.

53. Zhang W, Liu R, Chen Y, Wang M, Du J. Crosstalk between Oxidative Stress and Exosomes. Oxidative Medicine and Cellular Longevity. 2022 Aug 30;2022:3553617.

54. Cheignon C, Tomas M, Bonnefont-Rousselot D, Faller P, Hureau C, Collin F. Oxidative stress and the amyloid beta peptide in Alzheimer’s disease. Redox Biol. 2018 Apr;14:450–64.

55. de Oliveira GAP, Silva JL. Alpha-synuclein stepwise aggregation reveals features of an early onset mutation in Parkinson’s disease. Commun Biol. 2019 Oct 11;2(1):1–13.

56. Huang B, Wang W, Bates M, Zhuang X. Three-dimensional super-resolution imaging by stochastic optical reconstruction microscopy. Science. 2008 Feb 8;319(5864):810–3.

57. Folch J, Lees M, Sloane Stanley GH. A simple method for the isolation and purification of total lipides from animal tissues. J Biol Chem. 1957 May;226(1):497–509.

58. Costes SV, Daelemans D, Cho EH, Dobbin Z, Pavlakis G, Lockett S. Automatic and quantitative measurement of protein-protein colocalization in live cells. Biophys J. 2004 Jun;86(6):3993– 4003.

59. Adler J, Parmryd I. Quantifying colocalization by correlation: The Pearson correlation coefficient is superior to the Mander’s overlap coefficient. Cytometry Part A. 2010;77A(8):733–42.

