## Supplemental Data 1 for "Stress-Induced Alteration of Small Extracellular Vesicles Drives Amyloid-Beta Sequestration and Exacerbates Alzheimer’s Disease Pathogenesis"

Supplementary Figure 1.

A TEM EVs

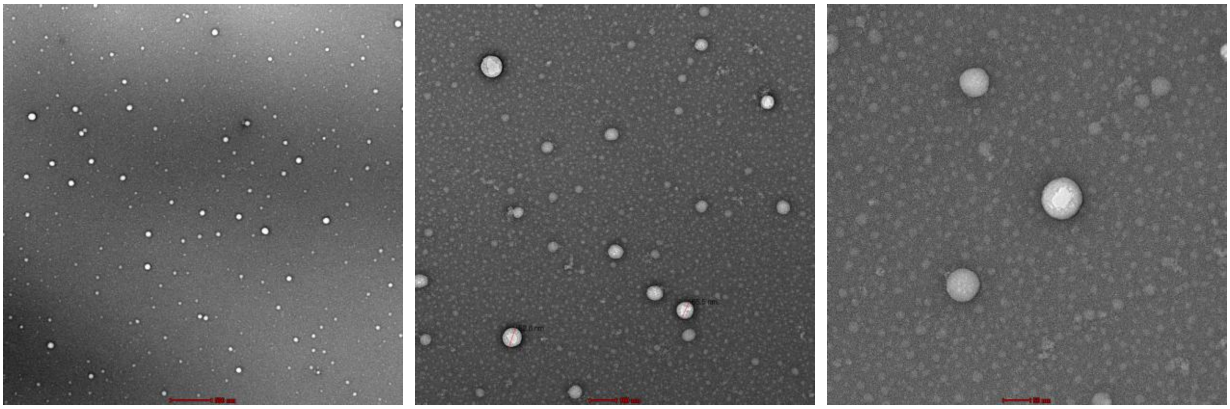

B Human CD9 Alexa Fluor® 488-conjugated Antibody tagged EVs

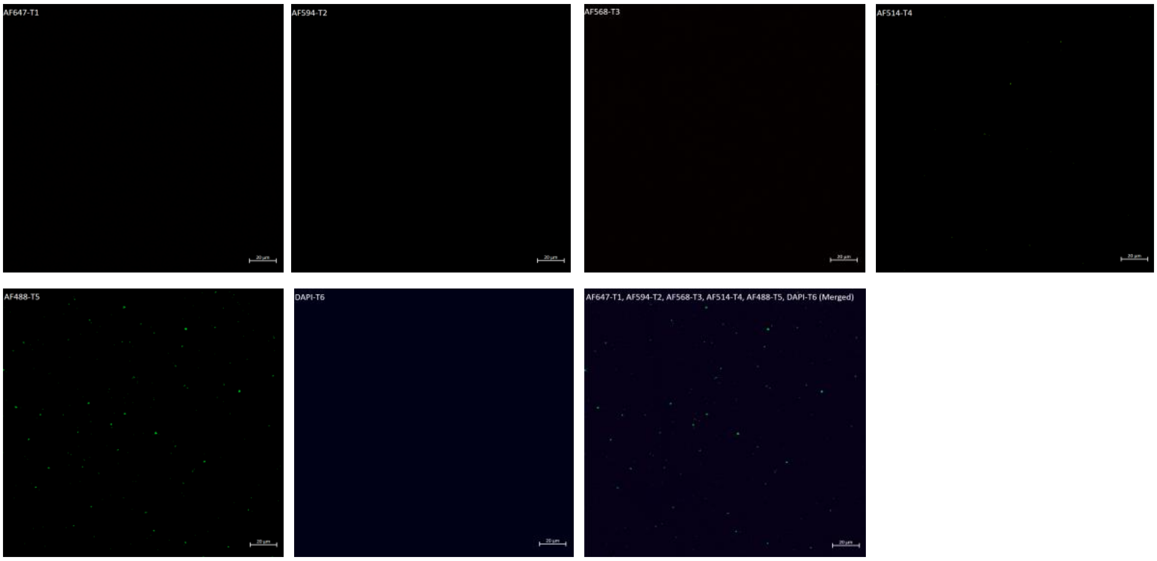


C Negative control

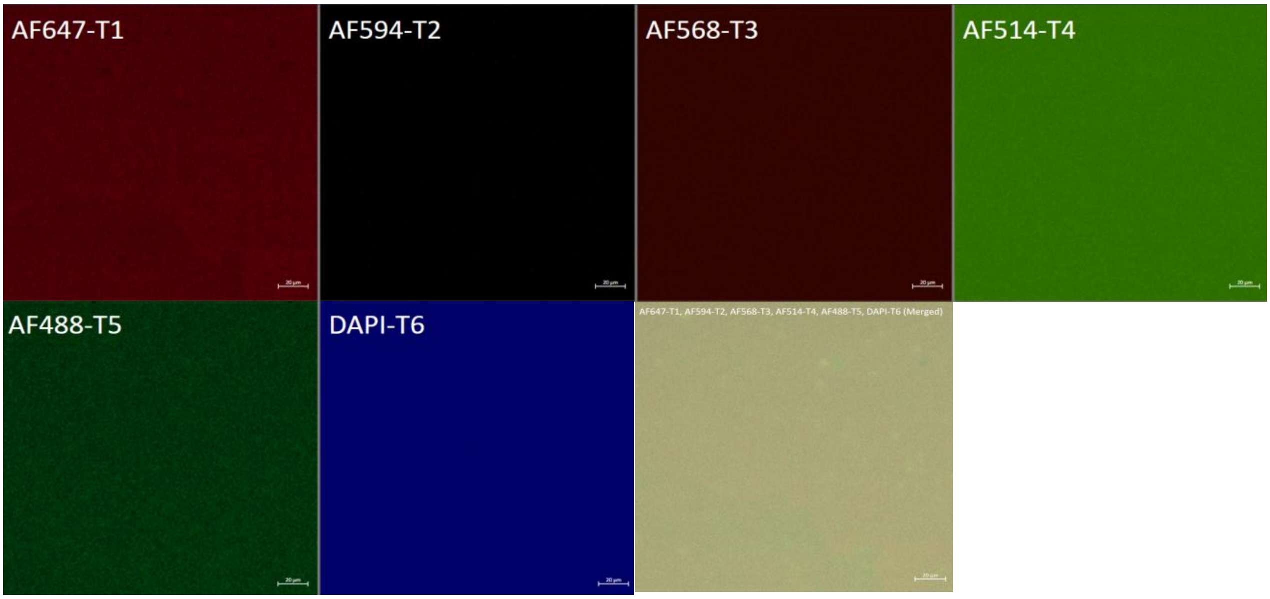

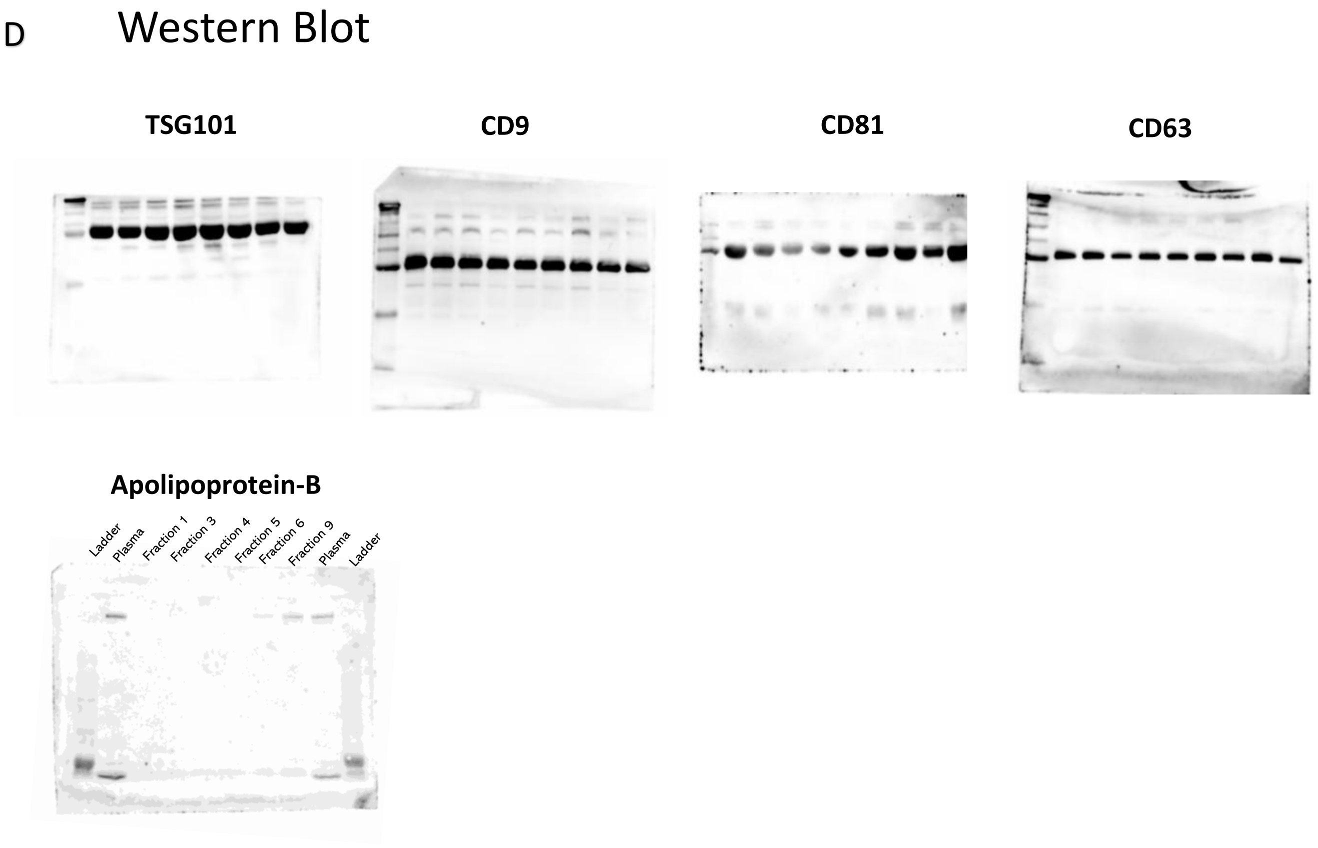


**Supplementary Figure 1:** TEM images of plasma-derived EVs (A); CFM Split image showing Alexa-Fluor-CD9 signals in all channels (B) and in negative control (PBS) (C); Western blot (Full blot) for EVs marker proteins and co isolated protein (Apo-B)(D).

Supplementary Figure 2.

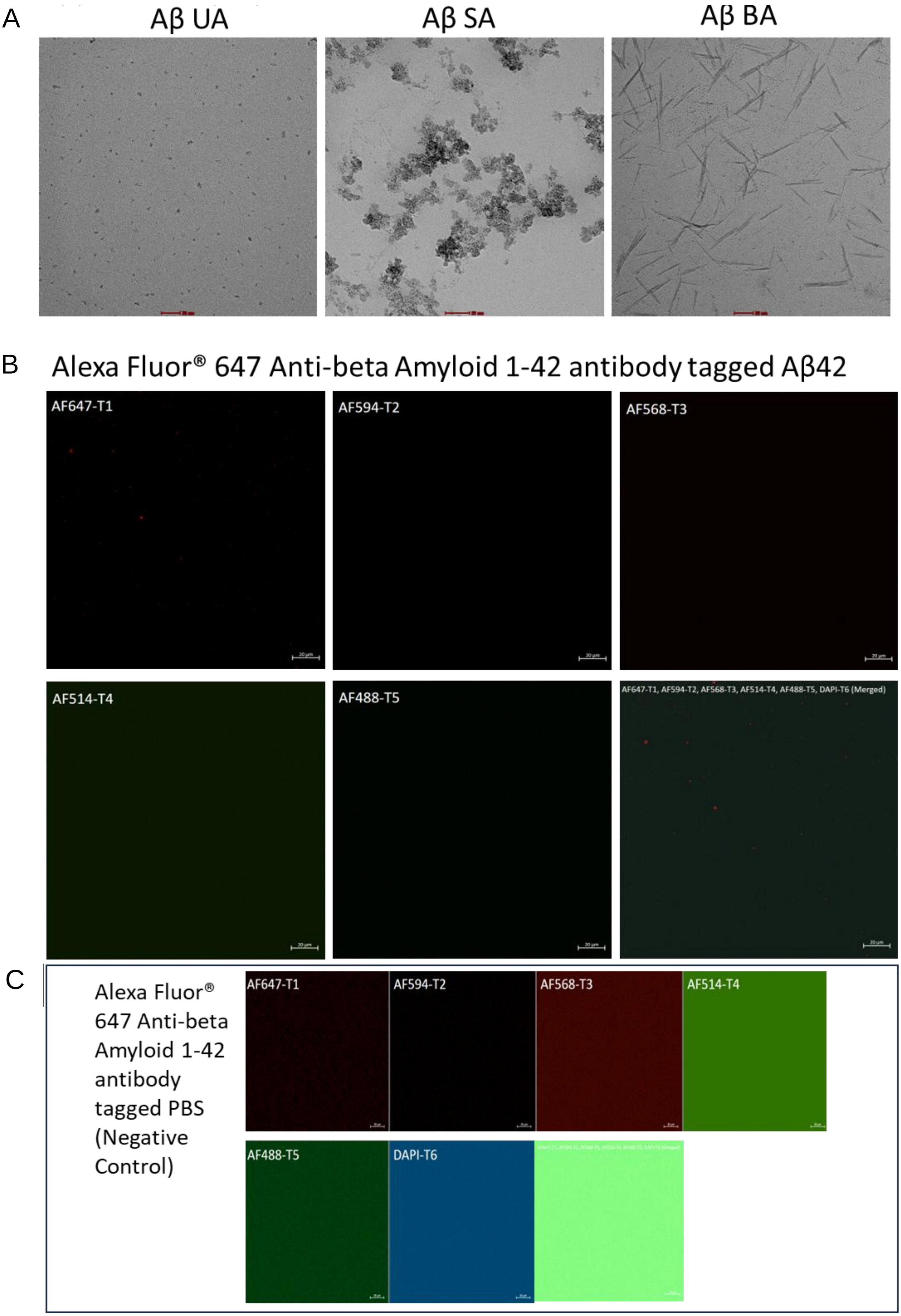

D

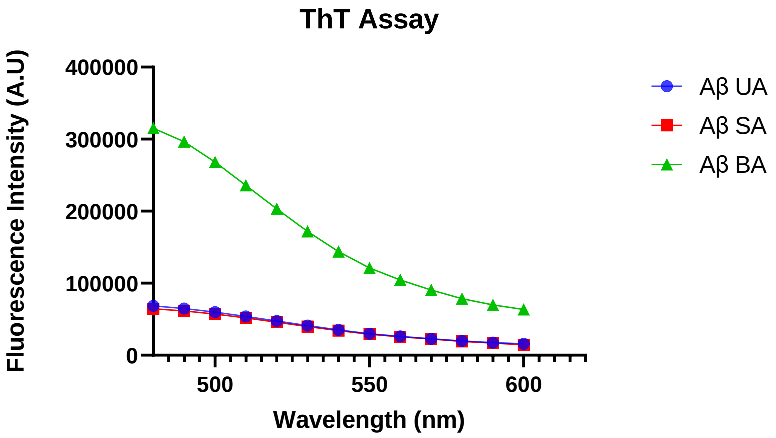


**Supplementary Figure 2:** TEM images of different Amyloid-β aggregates(A); CFM Split image showing Alexa-Fluor-647 Amyloid-β signals in all channels (B)and in negative control (PBS) (C); ThT Fluorescence for three Aβ groups (D).

Supplementary Figure 3.

A

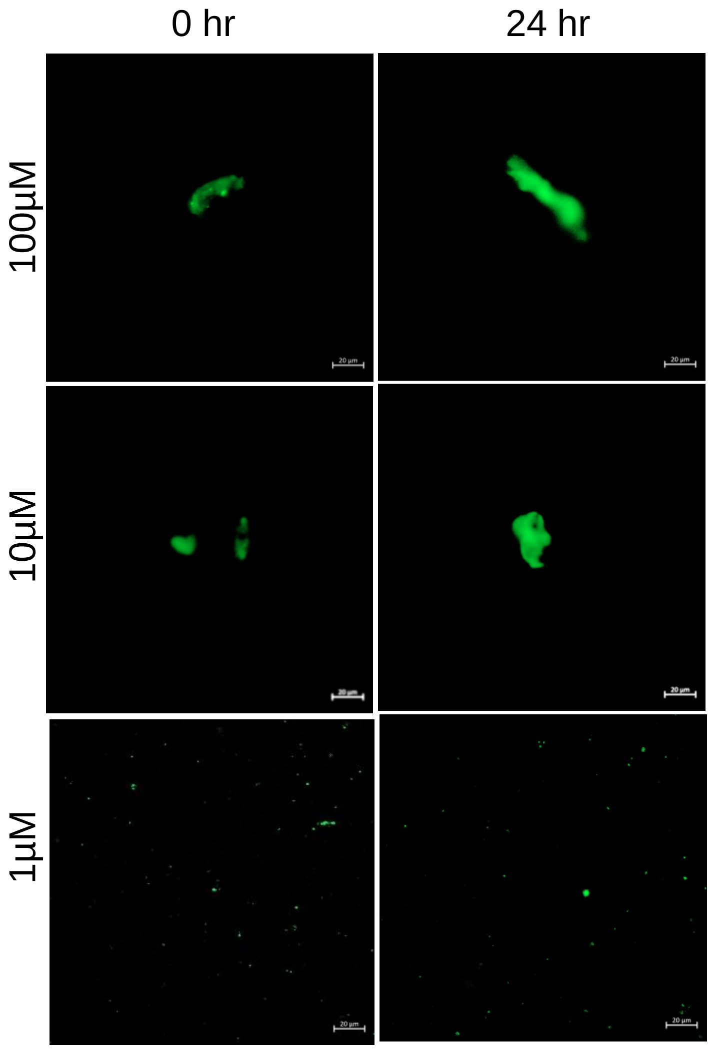


B

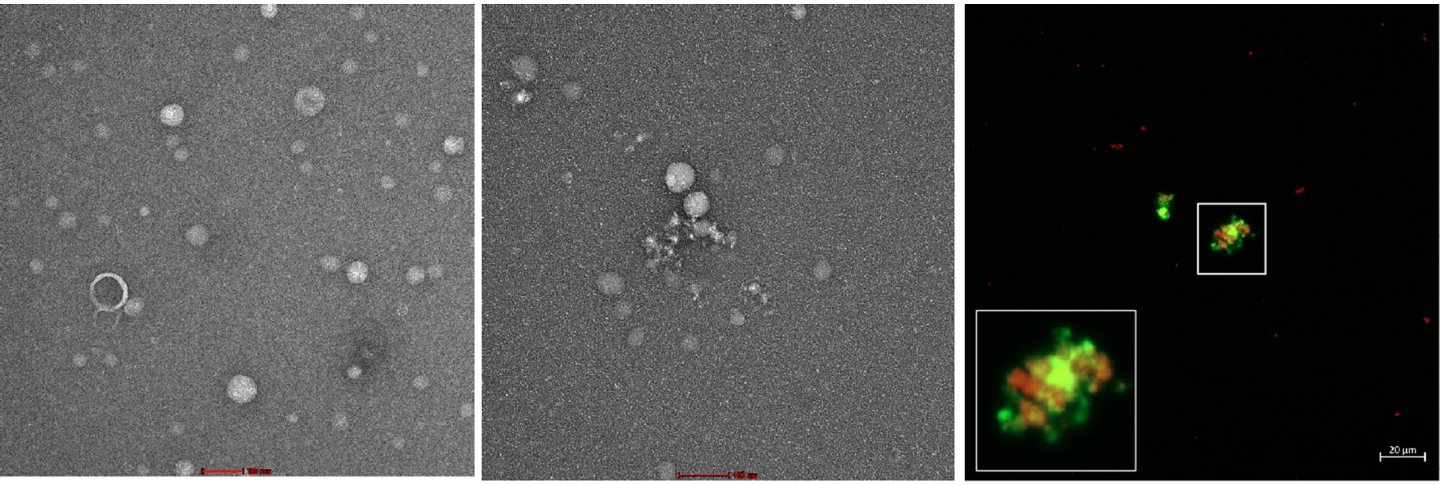

C

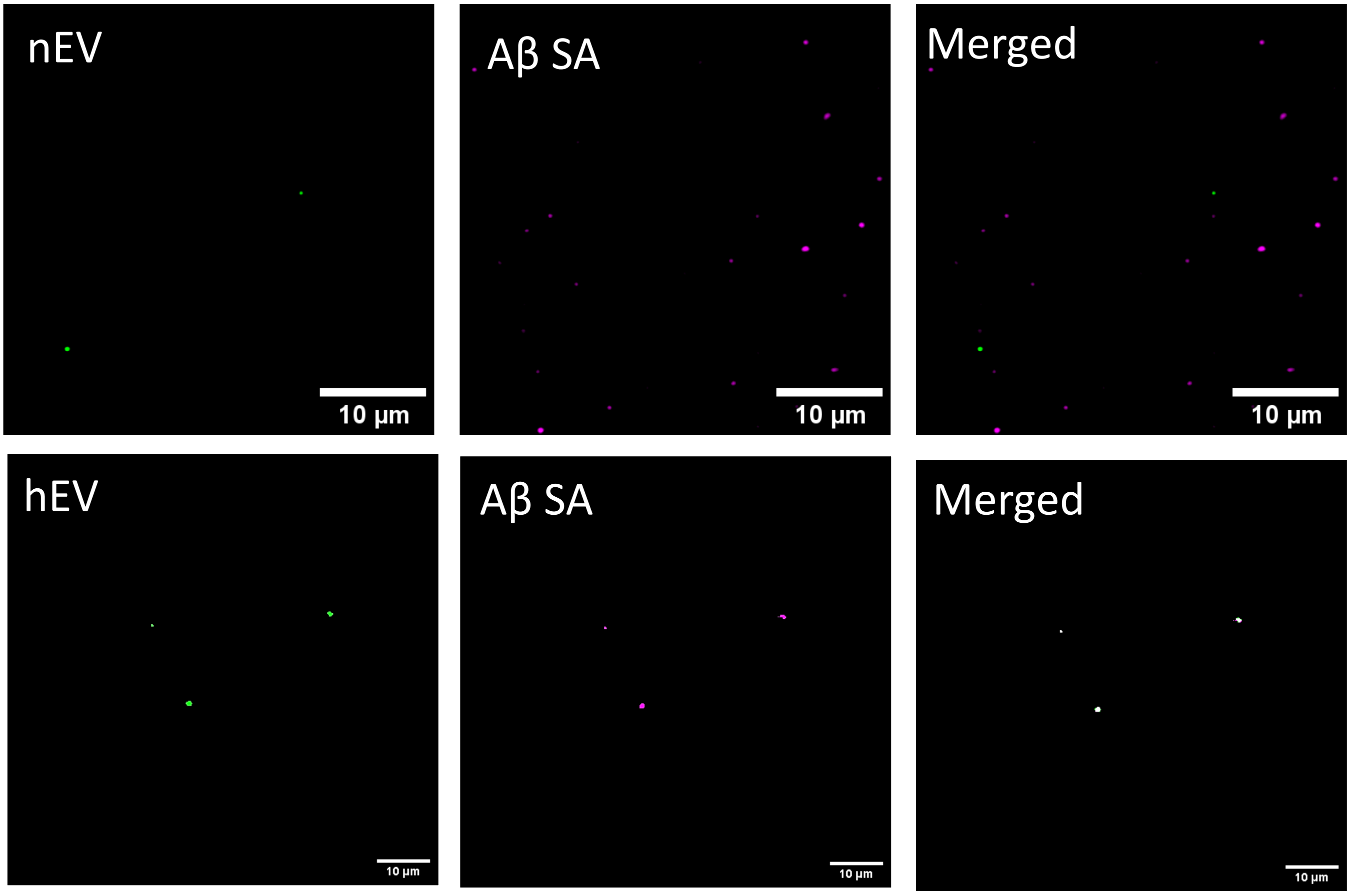
nEV
10 µm
Aβ SA
10 µm
Merged
10 µm
hEV
10 µm
Aβ SA
10 µm
Merged
10 µm

D

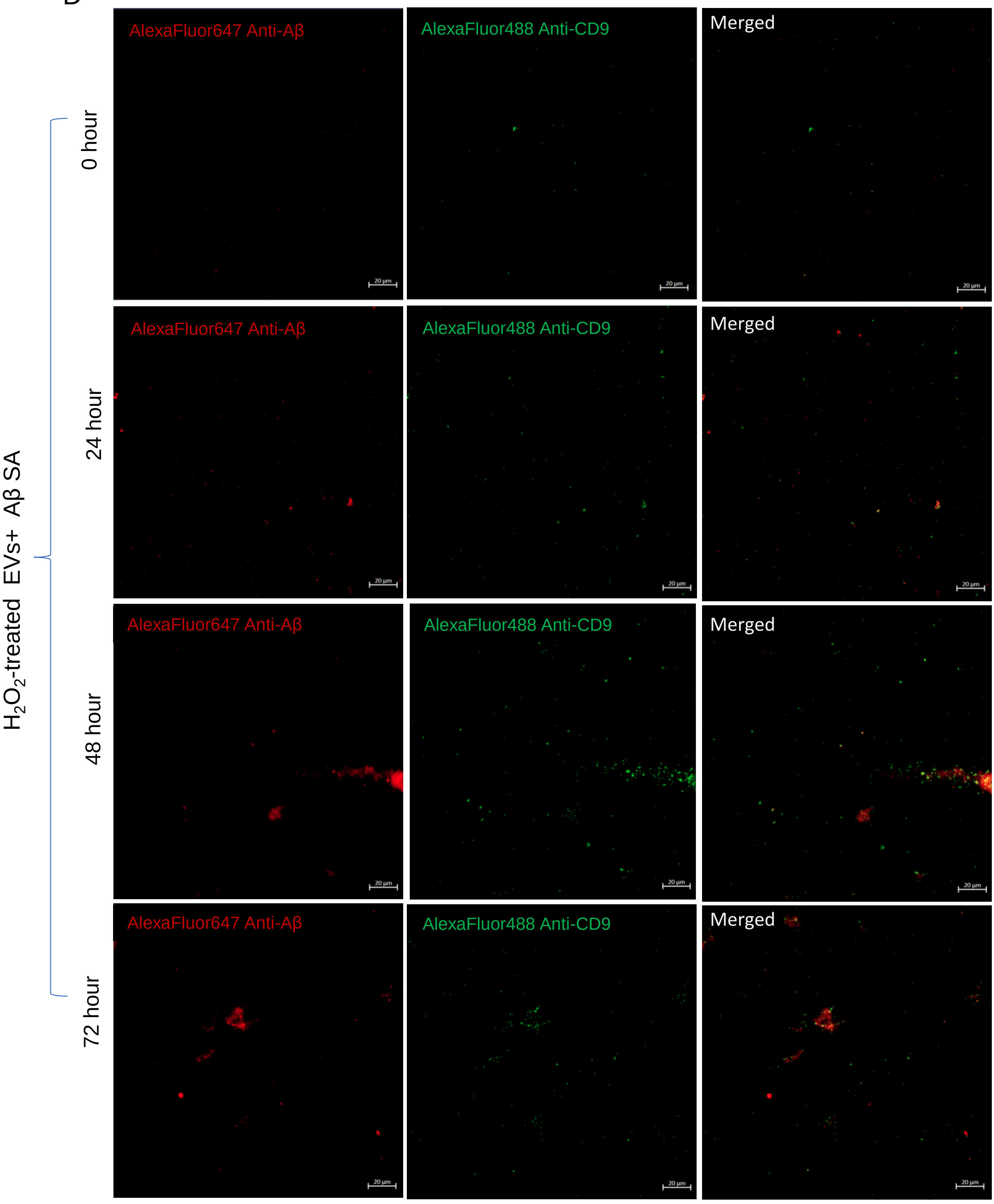

AlexaFluor647 Anti-Aβ
AlexaFluor488 Anti-CD9
Merged
0 hour
24 hour
48 hour
72 hour
H2O2-treated EVs+ Aβ SA
20 μm

E

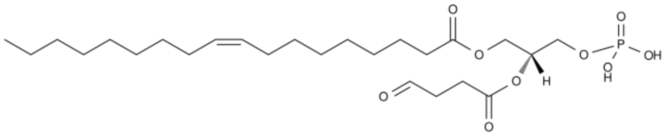

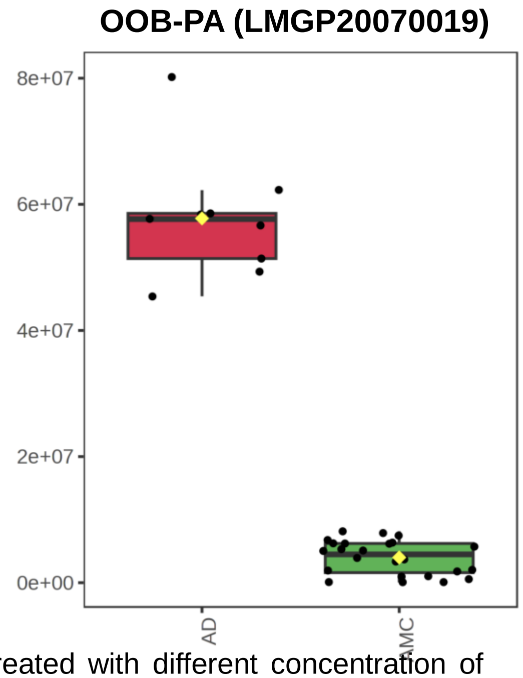


Category: Glycerophospholipid,
Class: Oxidized Glycerophospholipid

**Supplementary Figure 3:** Confocal image of EVs treated with different concentration of $H_2O_2$(A), Ultrasonicated Evs & Aβ(SA) (B), TIRF split images showing Alexa-Fluor-488 CD9 (Green) and Alexa-Fluor-647 Amyloid-β (Magenta) signals of: normal EVs+ Aβ(SA) & $H_2O_2$-treated EVs + AβSA. Colour white is the merged signal of EVs (Green) and Aβ (Magenta) (C), $H_2O_2$-treated EVs+ AβSA at 0-72hrs at 4℃. Colour yellow is the merged signal of EVs (Green) and Aβ (Red)(D)Scale bar= 20μm.(C) Elevated level of oxidized Glycerophospholipid in AD patients EVs (Unpublished data) (D).

### Supplementary Figure 4

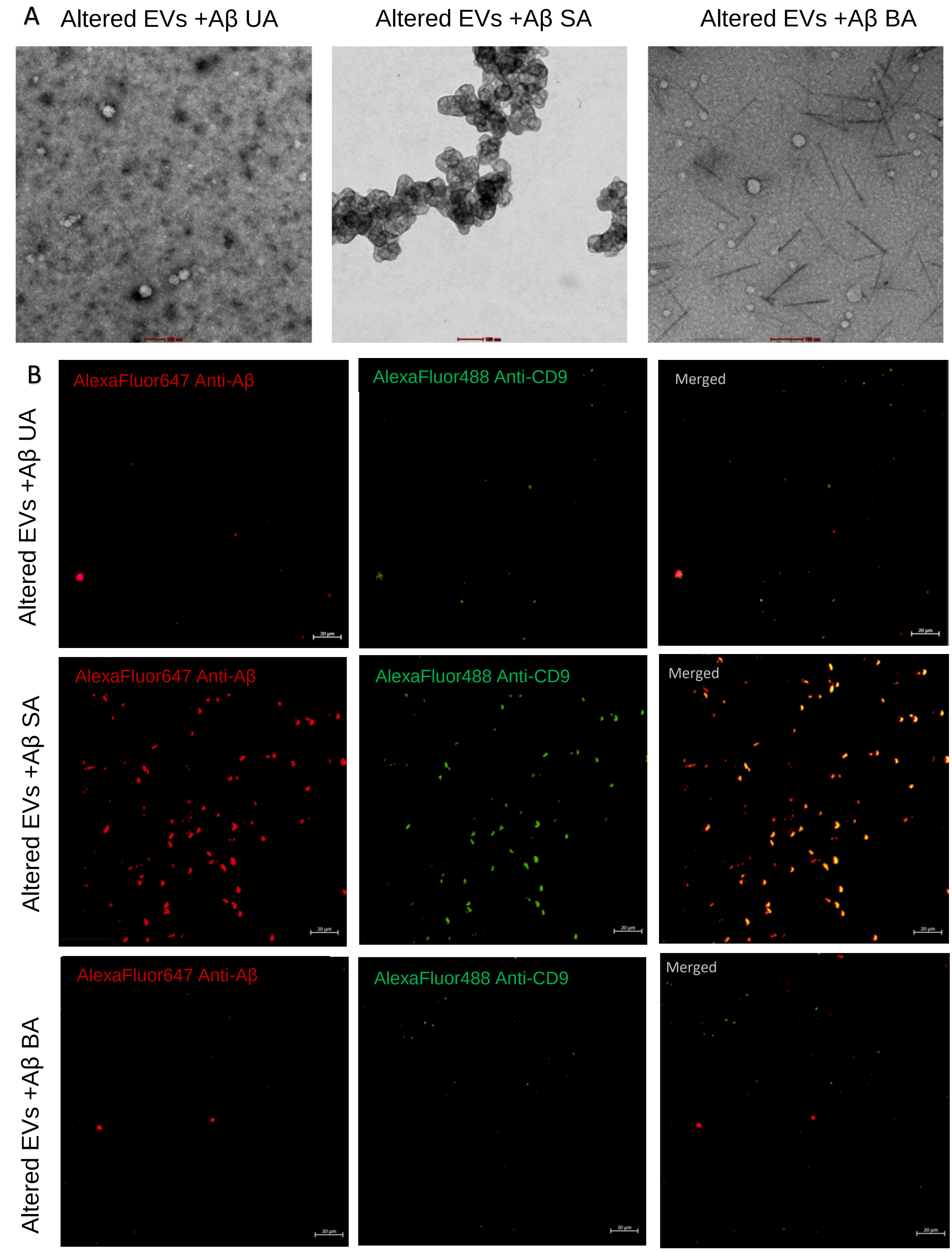


**Supplementary Figure 4:** TEM images of EVs and Aβ UA, SA and BA. Scale bar 50nm. (A); CFM Split image showing Alexa-Fluor-488 CD9 and Alexa-Fluor-647 Amyloid-β signals for all 3 groups. Colour yellow is the merged signal of EVs (Green) and Aβ (red). Scaler bar= 20µm.

Supplementary Figure 5

A

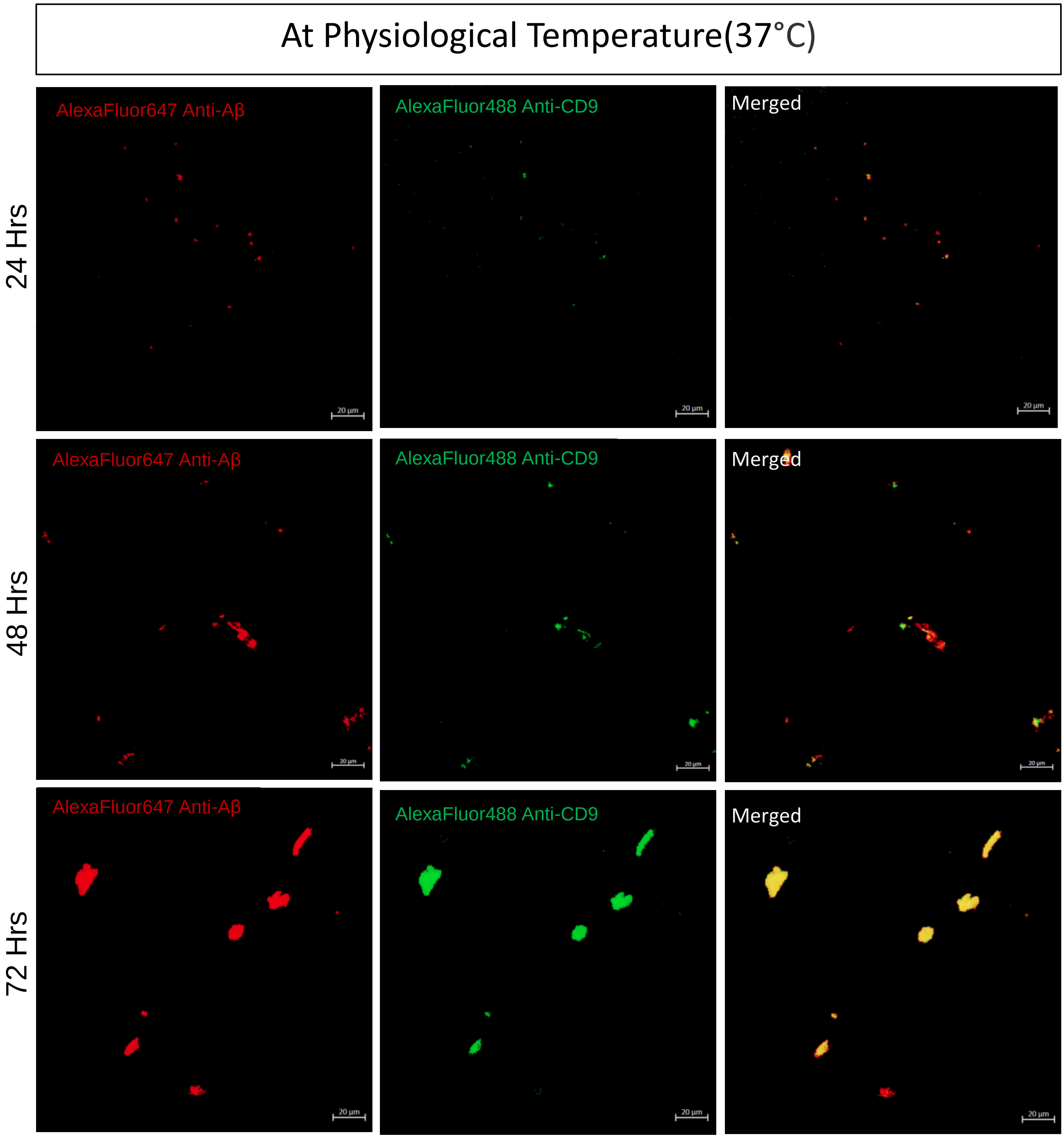

B

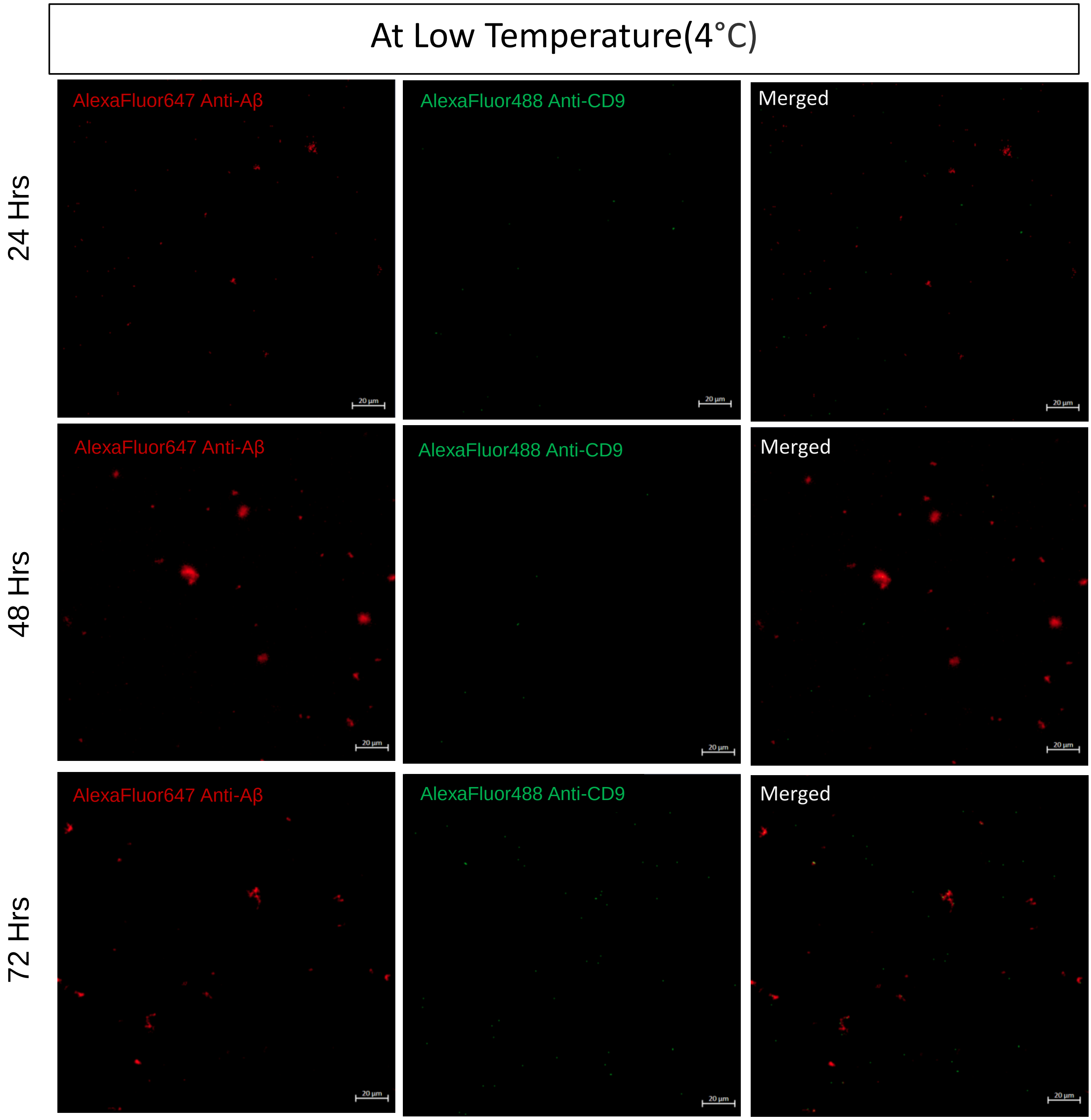


**Supplementary Figure 5:** CFM Split image showing Alexa-Fluor-488 CD9 and Alexa-Fluor-647 Amyloid-β signals of experimental group 24, 48 and 72 hours at 37°C (A) and at 4°C (B). Colour yellow is the merged signal of EVs (Green) and Aβ (red). Scaler bar= 20µm.

### Supplementary Figure 6

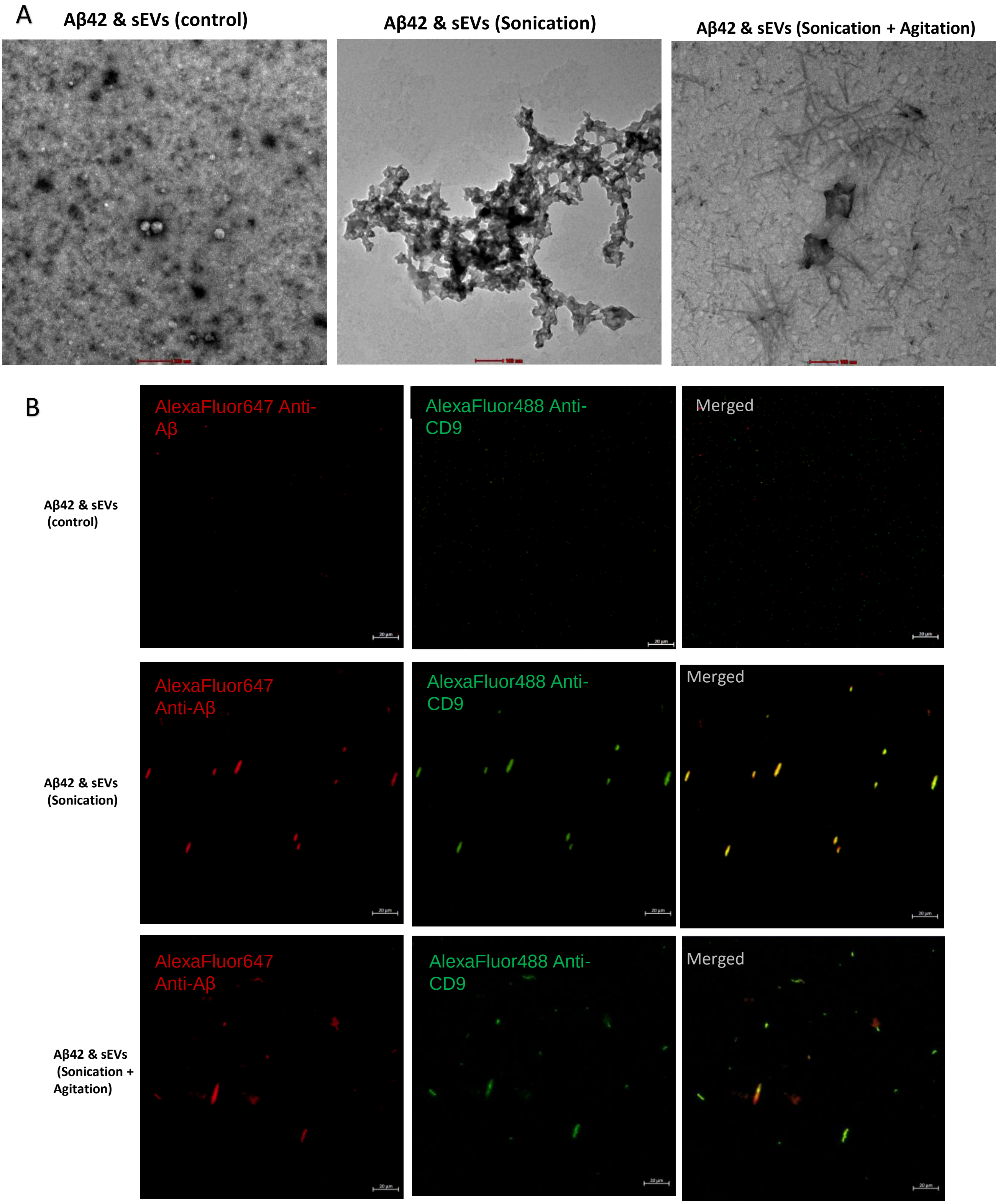

C

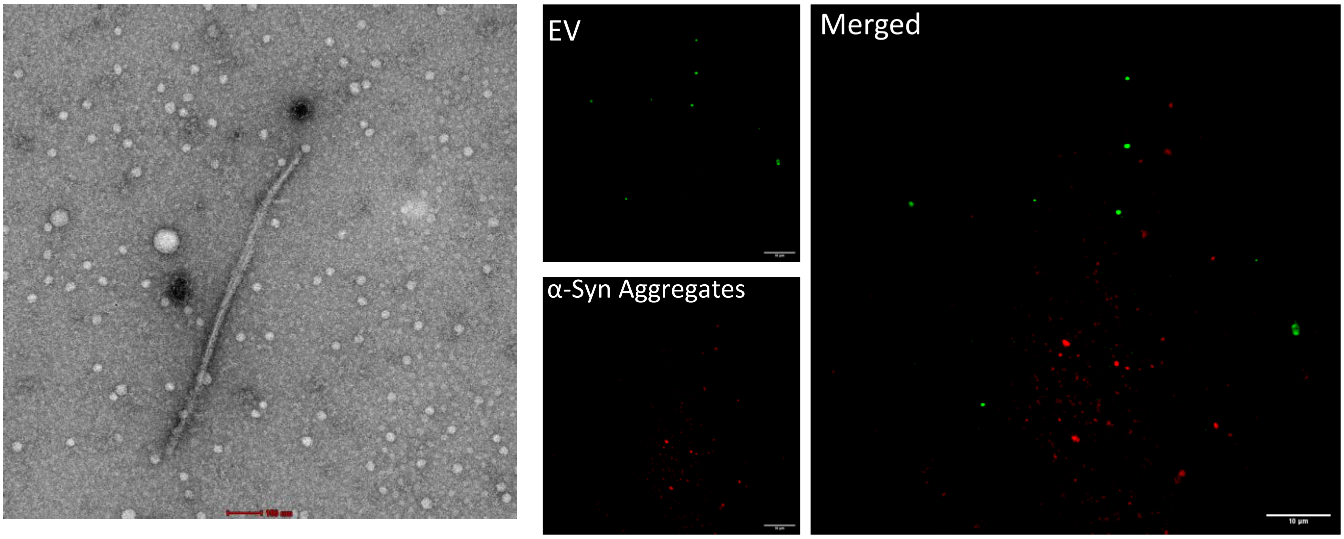


**Supplementary Figure 6:** TEM images of EVs and Aβ in control, sonicated and sonicated & agitated condition (Scale bar 50nm) (A); CFM Split image showing Alexa-Fluor-488 CD9 and Alexa-Fluor-647 Amyloid-β signals for all 3 groups (B). Colour yellow is the merged signal of EVs (Green) and Aβ (red). Scaler bar= 20μm. In-Vitro analysis of alpha-synuclein and EV (D)

### Supplementary Figure 7

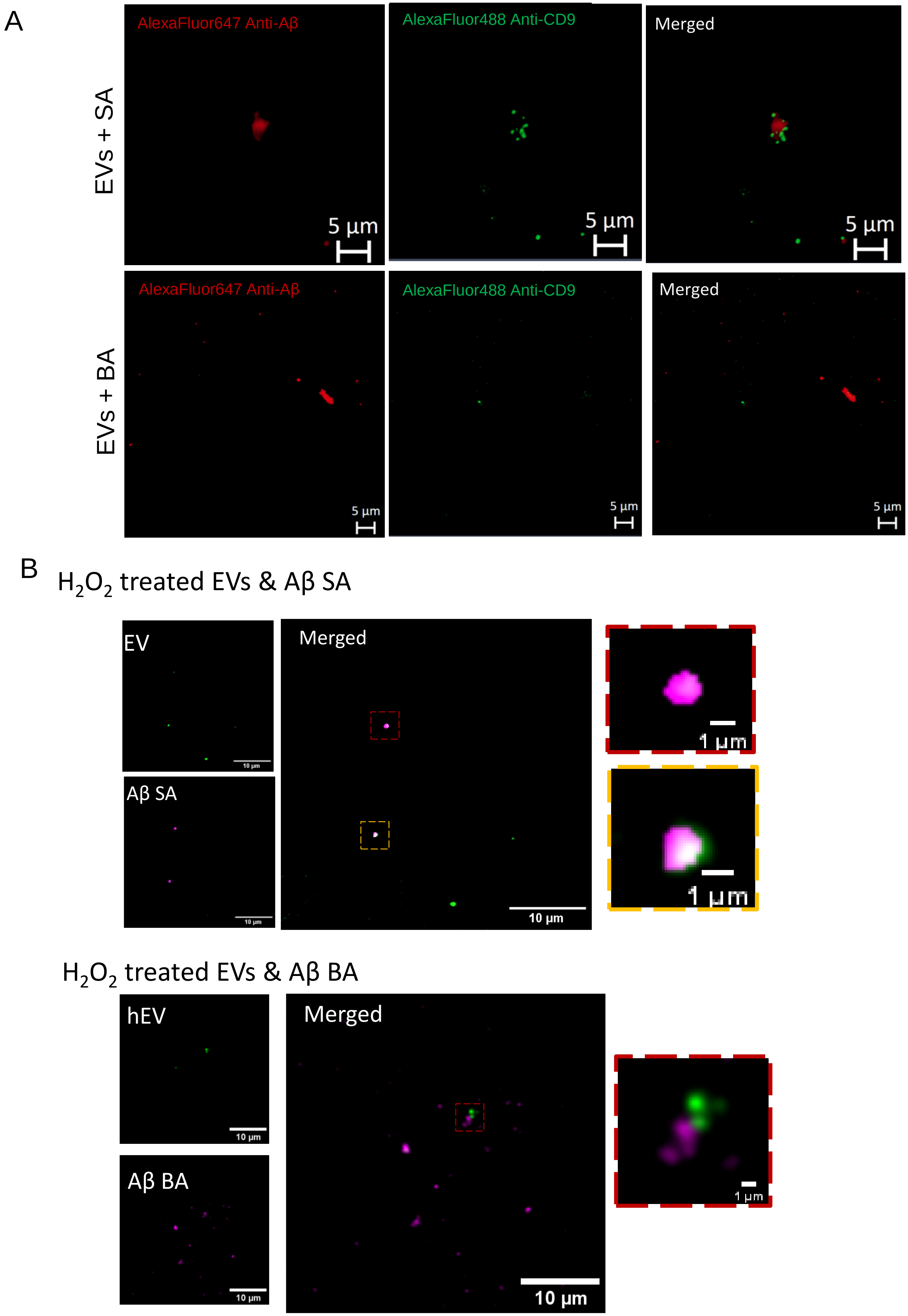

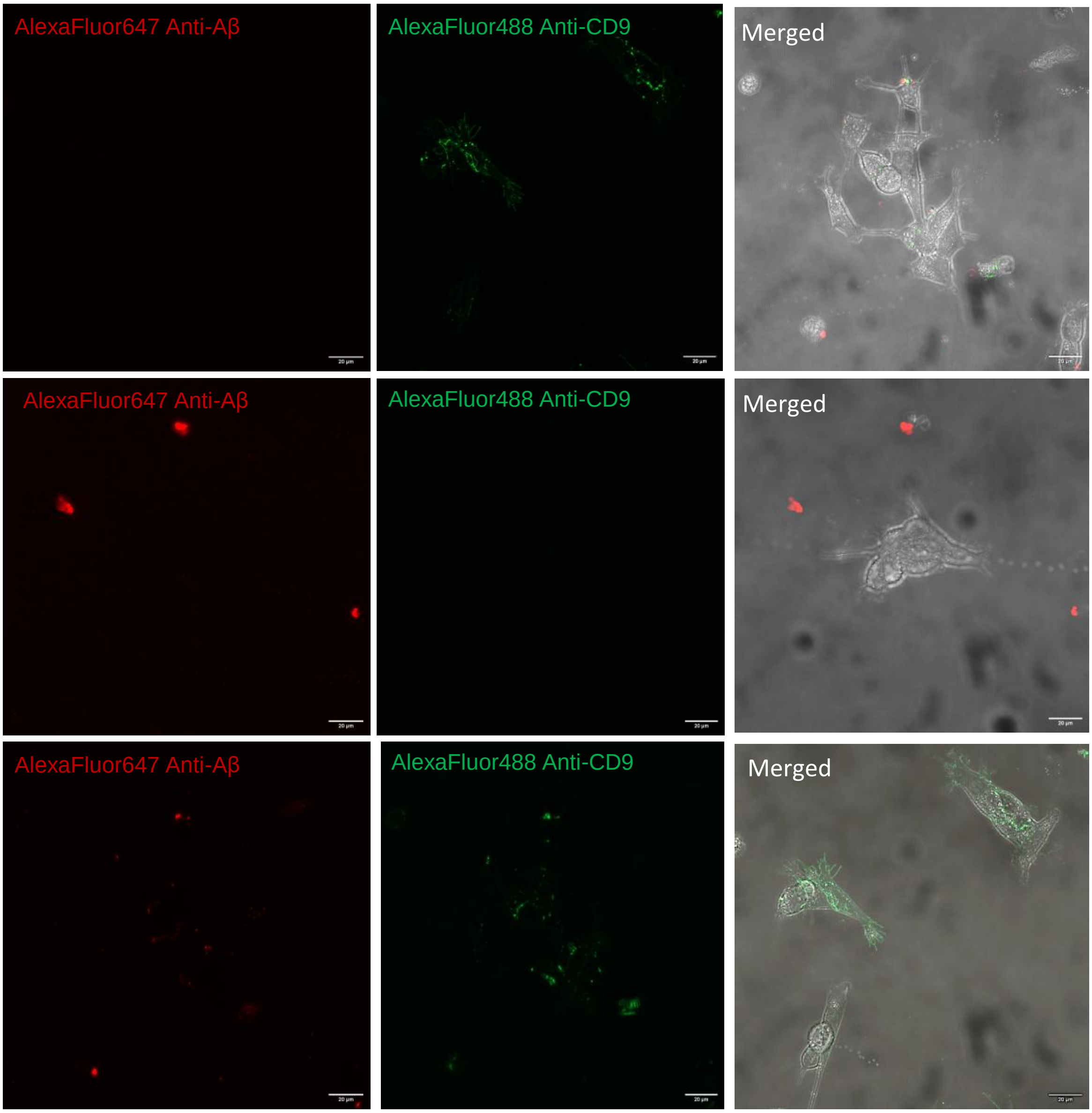


**Supplementary Figure 7:** CFM Split image showing Alexa-Fluor-488 CD9 (Green) and Alexa-Fluor-647 Amyloid-β (Red) signals of EVs+ Aβ SA and EVs+ BA under higher magnification shows EVs (Green) sequestering Aβ (Red). (A) CFM Split image showing Cell internalization of Alexa-Fluor-488 CD9 (Green) and Alexa-Fluor-647Amyloid-β (Red) signals of: EVs only; Aβ; and EVs and Aβ together (B). Scaler bar= 20μm.

### Supplementary Figure 8

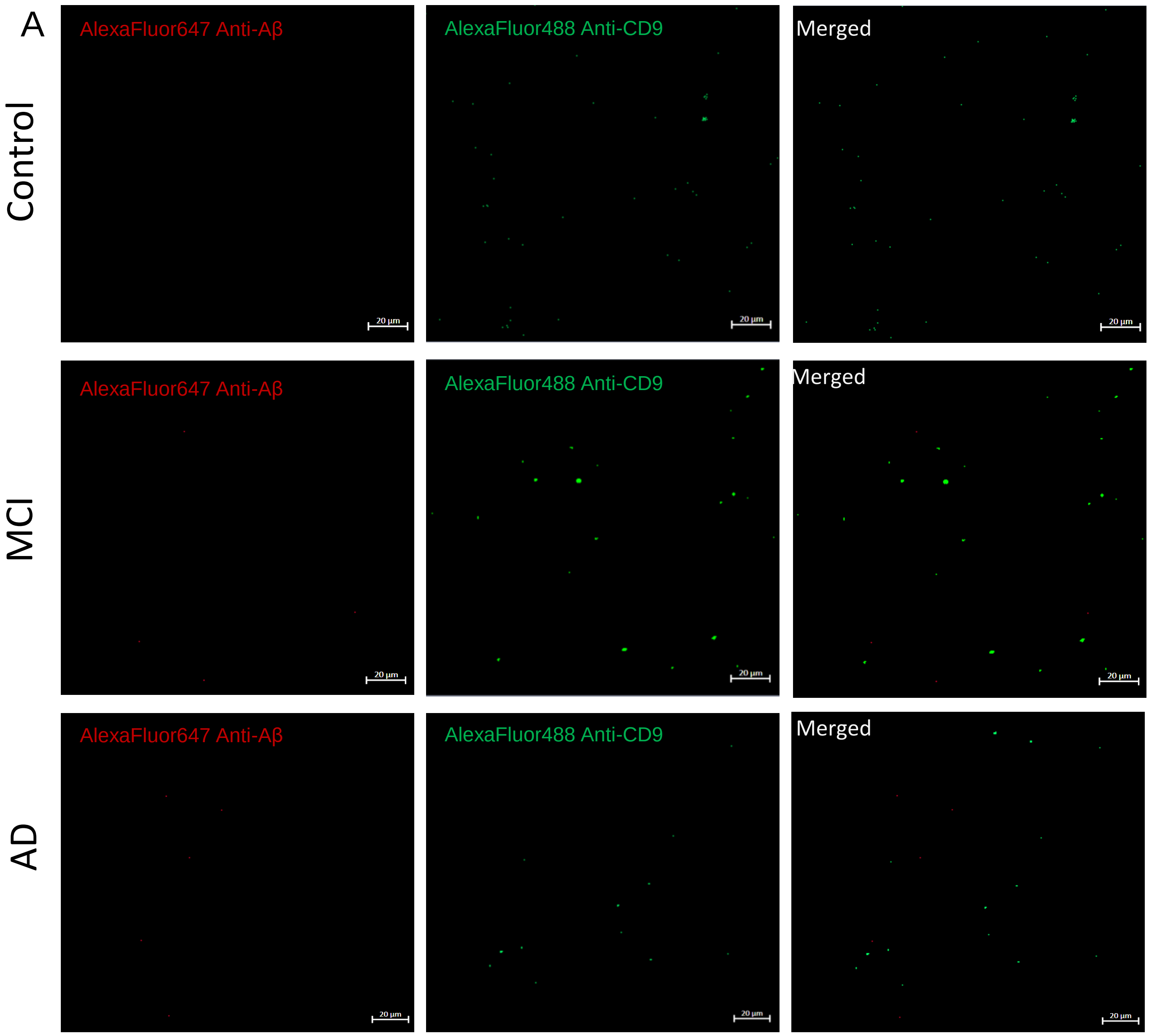

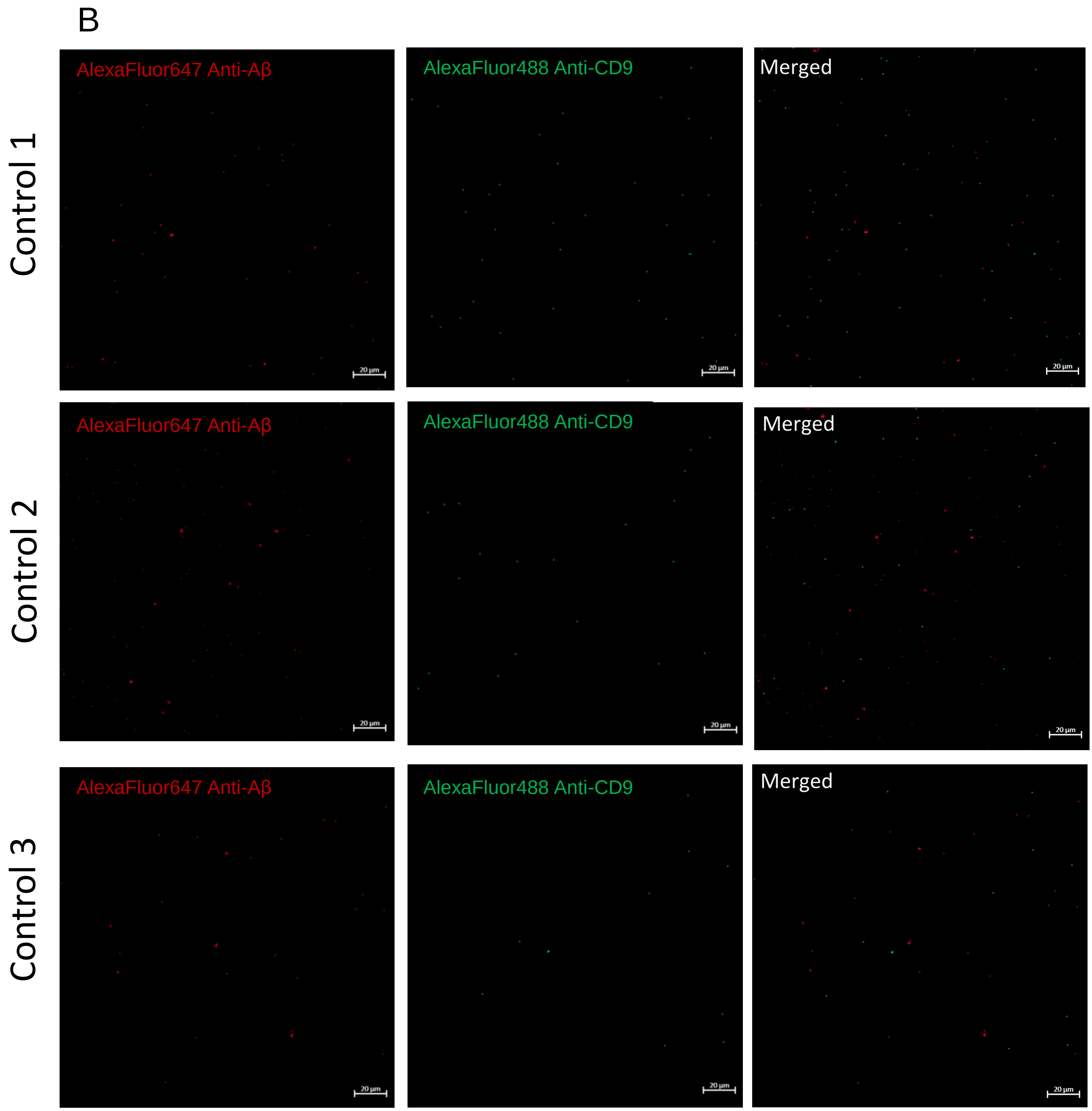
B
Control 1
Control 2
Control 3
AlexaFluor647 Anti-Aβ
AlexaFluor488 Anti-CD9
Merged
20 µm

C

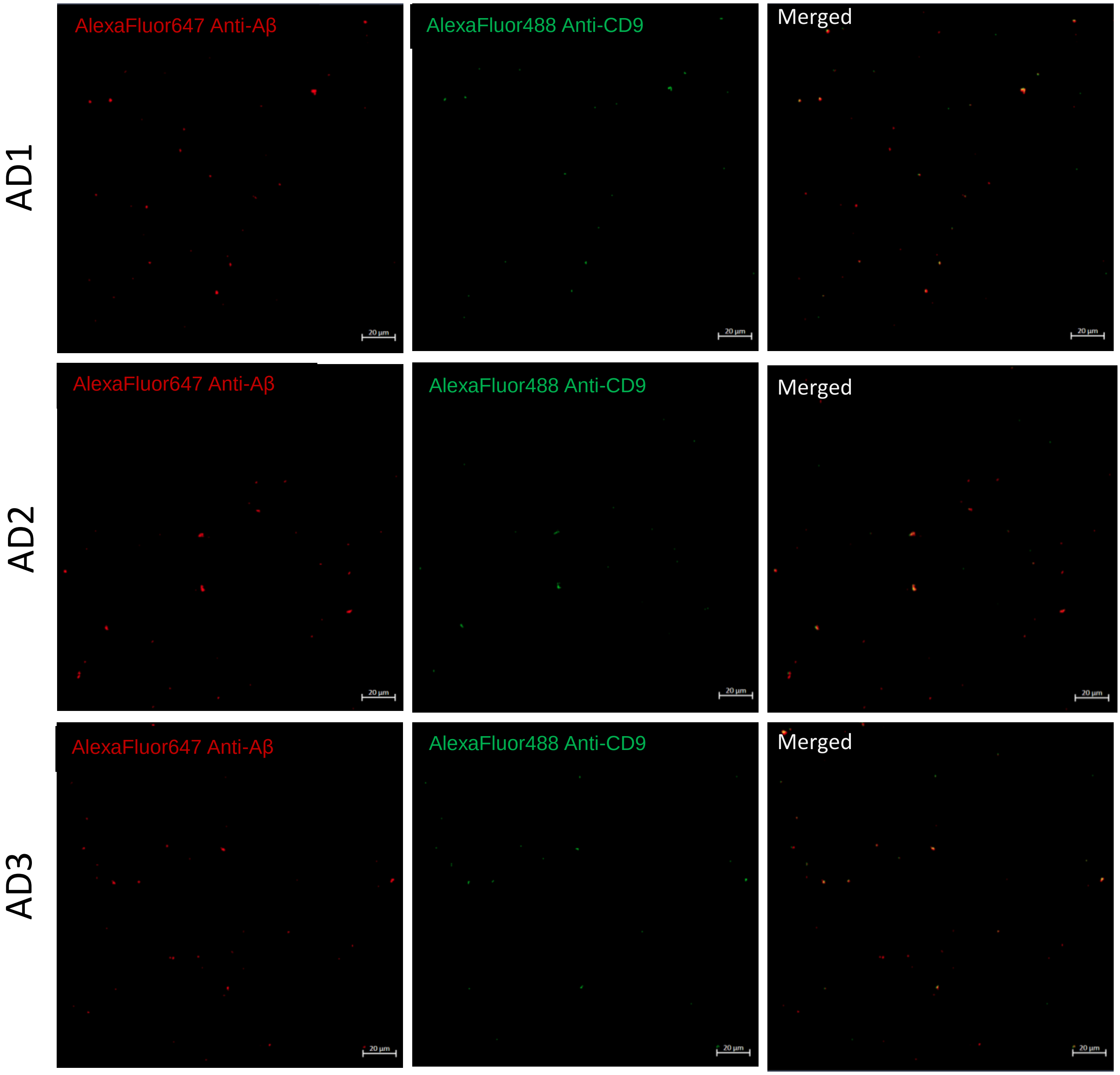


**Supplementary Figure 8:** CFM Split image showing Alexa-Fluor-488 CD9 (Green) and Alexa-Fluor-647Amyloid-β (Red) signals of: Circulating EVs in Age-matched control, MCI and AD patient (A); EVs from healthy control + Aβ SA (B); EVs from AD patient + Aβ SA (C). Scaler bar= 20µm.
